# 2D representations of DNA sequence show that most transversions are misaligned nucleotides associated with replication slippage

**DOI:** 10.1101/2024.07.17.603925

**Authors:** Albert J. Erives

## Abstract

Homologous sequences diverge in length via insertions and deletions (indels). Consequently, evolutionary genetic analyses routinely use methods to produce gapped alignment (GA). In GA, artificial null characters (gaps) are inserted into sequences so that nucleotide characters may be placed into homological correspondence within an alignment column. However, this approach sacrifices the homological correspondence of nucleotides diverging via tandem repeats (TRs). To address this deficit, we generalize GA with *micro-paralogical gapped alignment* (MPGA). While GA operates under a strict two-state homology model of *one-to-one* and *one-to-none* (i.e. one-to-gap) relationships, MPGA adds *one-to-many*, *many-to-many*, and *many-to-none* relationships. This expanded, multi-state homology model is motivated by DNA replication slippage (RS). RS produces short tandem repeats, constituting interrelated micro-paralogous sequences. Together, RS and TR-associated instability have a synergistic effect in the production of indels, which generate the need for gap insertions. MPGA reduces the computational cost of determining optimal gap insertions by reducing the number of gaps required by two-dimensional (2D) representations of sequence. A 2D representation of one sequence is achieved when tandem repeats are contracted into the same columns (dimension one) by occupying multiple rows (dimension two), an internal micro-paralogical dimension. To demonstrate the benefits and challenges of 2D representation, we develop a program called *LINEUP* and identify a pervasive fractal dimension in evolving sequences. We then demonstrate how *LINEUP*-generated 2D representations provide improved measures of substitution rates and transition-to-transversion ratios. Altogether, these results showcase significant new perspectives on basic mutational and evolutionary processes when multi-state homology models are adopted.

## Introduction

The evolution of a gene locus is driven by distinct molecular mechanisms that change its DNA sequence. These changes include: **(*i*)** nucleotide substitutions, **(*ii*)** short insertions and deletions (indels) <100 bp; **(*iii*)** gene conversions at small (<100 bp) to larger scales (>1kb); **(*iv*)** insertions and excisions of mobile elements, and **(*v*)** structural rearrangements such as inversions and translocations. These types of changes are detected by aligning homologous sequences. Sequence alignment for phylogenetic inference is geared toward producing “gapped alignment” (GA) as output. In GA, nucleotides in homologous sequences are aligned either to each other in the same alignment column or to gaps represented by null characters (typically dashes) inserted into homologous sequences. These two possibilities denote a two-state homological model whereby a nucleotide is either in a *one-to-one* relationship with a nucleotide in a homologous sequence or a “*one-to-none*” relationship with a gap. GA’s homological model incurs a cost with the computational complexity of determining the optimal placement of gap insertions. For this reason for example, the generalized affine gap penalty *p = g + m*(*x* − 1) can be used to limit the evaluation of alignment possibilities with an initial gap insertion penalty *g* and an additional gap extension penalty *m* scaled by *x* − *1*, where *x* is the total gap length (Gotoh 1982).

As GA is based on a two-state homological model (*one-to-one*, *one-to-none*), it is not predisposed to consider the likelihood that short indels originate from replication slippage (RS). RS is caused by the denaturation of a leading replication edge and the resumption of replication at a short offset from its prior position (Canceill and Ehrlich 1996; Canceill, et al. 1999; Viguera, et al. 2001). RS is responsible for generating the majority of short (<100 bp) tandem repeats (TRs). The probability of RS and instability is exacerbated by TRs themselves and increases logarithmically with repeat copy number (Ananda, et al. 2013). Maximum RS instability plateaus after about ten copies for all dinucleotides, trinucleotides, and tetranucleotide repeats, and a bit longer than ten bp for mononucleotide runs (Ananda, et al. 2013). Thus, RS-mediated indels and their inherent TR biology are significant drivers of length divergence in homologous sequences.

Because of the causal interconnectedness of RS, TRs, and indels, a more accurate and evolutionary-framed approach to nucleotide sequence alignment might be achievable with an expanded multi-state homological model that also includes *one-to-many*, *many-to-many*, and *many-to-none* relationships between nucleotides. Thus, this study is concerned with gathering any and all insights into the feasibility of 2D representations. Furthermore, it was not initially obvious whether such representations were generally even possible for all sequences.

I refer to the expanded multi-state homological approach to GA as micro-paralogical gapped alignment (MPGA > GA). MPGA generalizes GA by representing sequences in a two-dimensional format with certain necessary constraints described here. The goal of 2D representations is achieved when tandem repeats in one sequence are placed into the same alignment columns (dimension one) by occupying multiple rows (dimension two), a micro-paralogical dimension. Here, I introduce a program called *LINEUP* to explore basic issues involved in 2D representation of a sequence (“self-alignment” of internal micro-paralogical content). Remarkably, 2D preparations of homologous sequences reduce the number of gaps required in their subsequent alignment. I then use *LINEUP* to identify dinucleotide repeat depletion and trinucleotide repeat enrichment of protein-coding sequences. Last, I build on this finding to show how the transversion mutational rate may be much lower and the insertion and deletion rate much higher than current estimates. For example, half of all apparent substitutions in the highly conserved *gapdh* gene are the result of canonical base pairing with adjacent sequence and are responsible for most apparent “transversions” seen in 1D alignments. These results demonstrate how the two-dimensional alignment of MPGA can provide new perspectives on evolutionary and mutational processes.

## Results

### Expanding the set of symbolic tokens used in GA for MPGA

With an expanded model of homology states that encompasses paralogical relationships, it is desirable to expand the set of symbolic tokens used in GA beyond the four nucleotides (**A**,**C**,**G**, and **T**) and the null character (“**-**”). Basic rules for tokenizing the inference of TRs in a proper 2D representation have been previously reported and can be used to ensure homological integrity of 2D representations (Erives 2019). Here, 2D representations are read from left to right, then from top to bottom. A replication slip character “**/**” is introduced to indicate where TR-like content continues in the next row at lower numbered columns. Dots or spacer characters (“.”) are used for clarity to highlight empty parts of a row before a sequence recontinues after its first row. Thus, spacer characters are used to correctly position the repeating units of a TR after the initial unit in the first row. Spacer characters should not be confused for null characters, which will continue to represent gaps per general convention. Last, terminal characters “**>**” facilitate reading where a sequence begins and ends, as the termini of one sequence may now occupy different rows.

The expanded set of tokens used in the 2D representations of MPGA is demonstrated below for the example of a CAG repeat expansion, in which a single CAG trinucleotide in one sequence has a *one-to-many* relationship with three copies in a homologous sequence, i.e. (CAG)_3_. In the 2D alignment below, the single CAG trinucleotide of sequence one is homologically related in a *one-to-many* relationship with three copies of the expanded CAG trinucleotide of sequence two. (The flanking n’s in the example below represent homologous anchor sequences conserved in both sequences.)

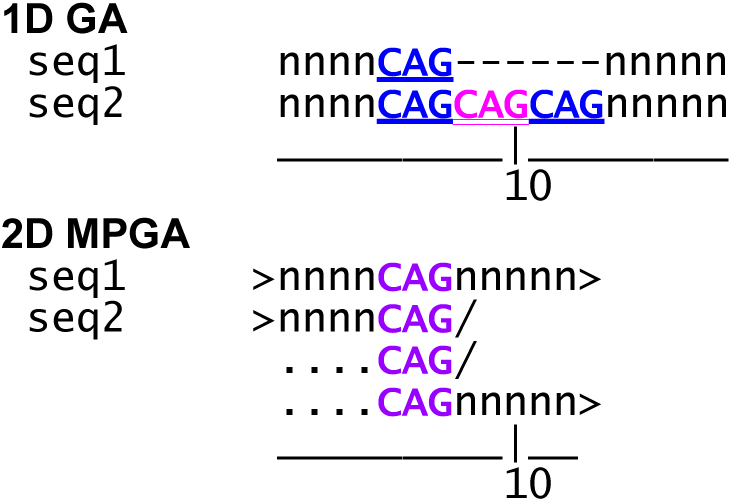

Note that in the above 2D MPGA, gaps were not required to successfully align a pair of homologous sequences of different lengths. In MPGA, gaps are inserted *after* the contraction of TR content only if they are still required.

2D alignments have widths shorter than 1D widths. I thus refer to the inference of replication slips in a 2D alignment as “*cinching*” the 2D width of a sequence (i.e., reducing the number of alignment columns). As will be shown, the cinching of the 2D widths of biological sequences results in the closing of many indel-associated gaps. As such, 2D cinching operations reduce the computational complexity of gap insertion.

### Exploring micro-paralogical self-alignment with the LINEUP program

To demonstrate the utility of 2D alignment and the *LINEUP* program, Figure 1 shows the outputs of multiple sequence alignment (MSA) of four sequences with divergent TR content (Fig. 1A, input) using various pure GA methods (Fig. 1B–G) versus the 2D MPGA of *LINEUP* (Fig. 1F). Only a 2D alignment can produce a gap-free 2D MSA, as achieved by *LINEUP*. This does not mean that 2D alignment does not use gaps; it only means that inferred replication slips of tandem repeats are invoked before the insertion of gaps. In the Fig. 1F example, 2D alignment does not need gaps to restore uniformity to the 2D widths of each sequence (24 bp). Furthermore, auto aligning each sequence to its internal repeat content is sufficient to render perfect alignment in this case.

**Figure 1.**
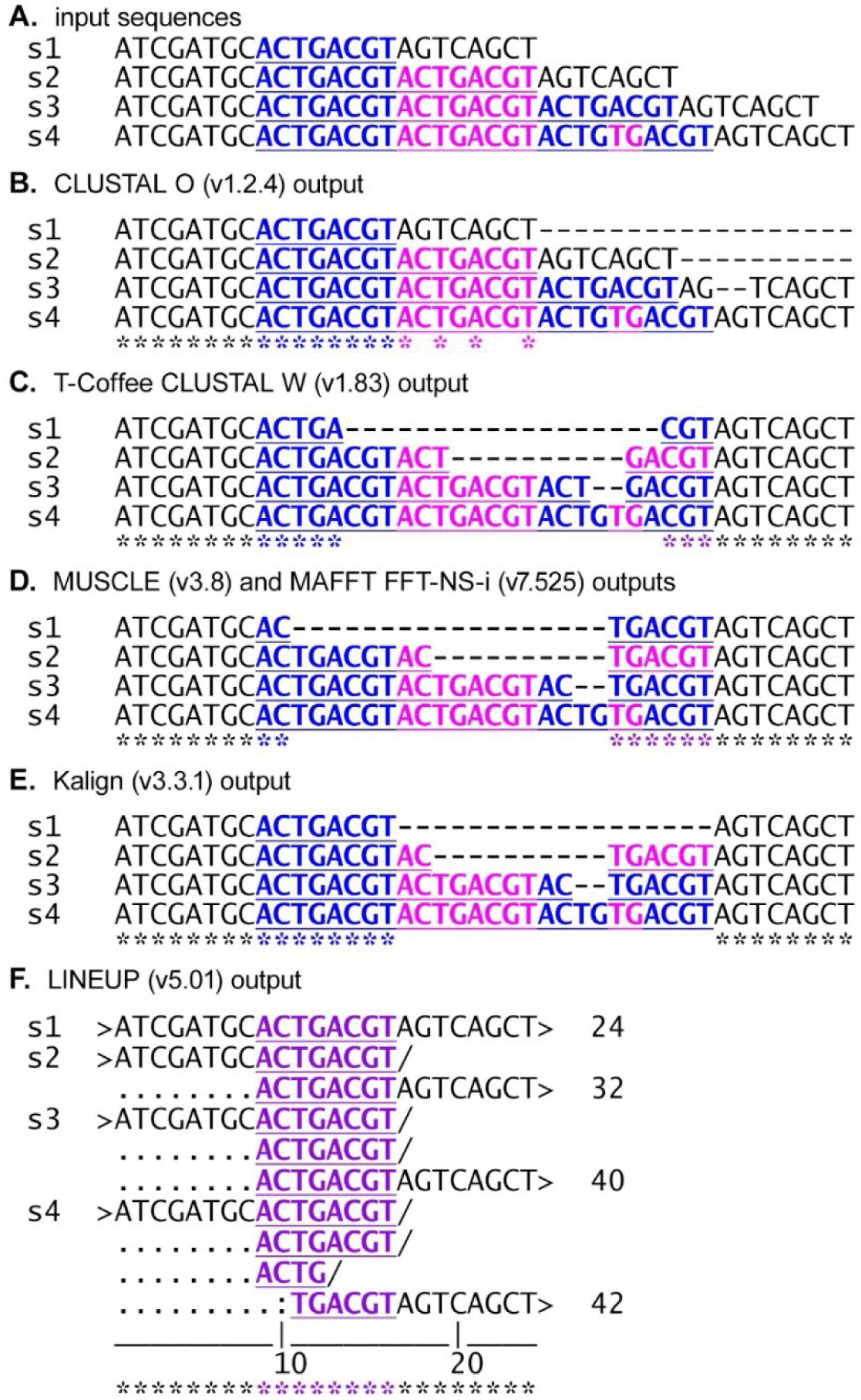
Standard GA algorithms are not designed to cinch TR content. (A) Shown are four homologous sequences (“s1” to “s4”) that differ in copy number of an 8-mer sequence (alternating blue and magenta sequences). Sequence s1 has a single (unrepeated) copy of the 8-mer sequence, while s2 has two copies. Sequences s3 and s4 have three copies each, but s4 has an additional dinucleotide “fractal” repeat within the third 8-mer parent unit. (B–E) Four different gapped alignment outputs are shown from five different MSA programs. These MSAs chop up repeat units in a TR-unaware manner (compare color-highlighted TRs), and only a subset of columns contains perfect nucleotide identity across all sequences (asterisks). (F). The resulting 2D MSA produced by *LINEUP*-generated self-alignments demonstrates how 2D self-alignment reduces the gap insertion problem. 2D alignment symbols: “>” marks the beginning and end of a sequence, which may occur in different rows; “/” marks the location of an inferred replication slip and the continuation of a sequence in the row below; “.” marks empty positions, not to be confused for null characters (“-”), which are reserved for indels.

Producing 2D self-alignment with perfect homological integrity of columns is only a superficially simple aim. Development of the *LINEUP* program revealed a profoundly rich world of TR biology and interesting problems inherent to 2D self-alignment. The foremost challenge is the impossibility of implementing a greedy approach whereby all tandem repeats are cinched in a 2D auto-alignment. The reason is that overlapping repeats often cannot be cinched simultaneously while preserving the homological integrity of columns. This is not an infrequent problem in DNA sequences because the instability of TRs frequently generates overlapping TRs with different *k*-mer sizes and/or related sequences. This challenge motivates the implementation of two algorithmic capabilities. The first capability is to use a suitable scoring system to decide between incompatible, overlapping TRs. The second capability is to evaluate the cinching or cycling frame of a repeat to find a frame compatible with overlapping repeats. All *k*-mer repeats ≥ 3*k*-1 in length will have *k* cycling frames. For example, 5′-<u>CAGCAGCA</u>T can be cinched as (CAG)_2_CAT, C(AGC)_2_AT, or CA(GCA)_2_T.

Figure 2A shows *LINEUP*’s output for a sequence containing multiple overlapping repeats with a choice of cinching or cycling frames. This example demonstrates that a greedy 2D alignment algorithm that cinches the maximum number of repeat units for each TR is not guaranteed to produce a non-pathological 2D alignment with homological integrity of columns. Pathological 2D alignments are defined as those with nucleotides accidentally placed into the same column as a secondary consequence of cinching in other columns. Thus, to ensure compatibility with the overlapping 4-mer TR, the same 5′-TGTGTG sequence in each of the four knots is sometimes cinched as (TG)_3_ (greedy), but sometimes less greedy, as T(GT)_2_G (Fig. 2A).

**Figure 2.**
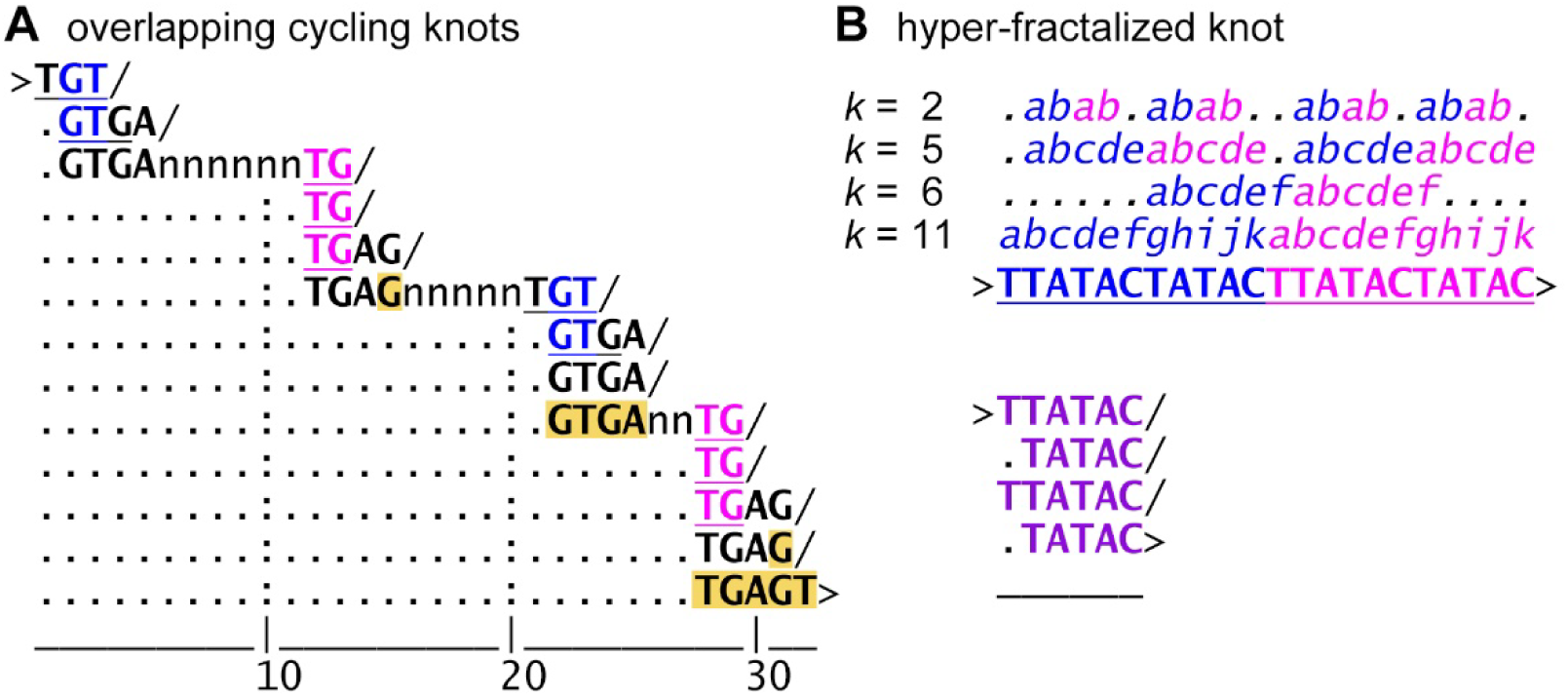
Two-dimensional self-alignment problems. (A) 2D self-alignment must identify compatible cinching frames for overlapping repeats, as shown for this 68 bp sequence that was optimally 2D-aligned by *LINEUP*. This sequence has four TR “knots” separated by n’s. The n’s represent ambiguous sequence, which *LINEUP* interprets as “un-cinchable.” Each of the four TR knots begins with the same 11 bp sequence. At each of the four knots, there are two overlapping TRs: a dinucleotide repeat (underlined) and a downstream tetranucleotide repeat. At each knot, the copy number of the *k*=4 repeat changes because of additional sequence (highlighted in yellow). This extra sequence causes *LINEUP* to choose different compatible repeat frames for the overlapping TRs to minimize the cinching width. For example, the dinucleotide repeat is cinched alternately as (GT)_2_ (blue) or (TG)_3_ (magenta). Similarly, different cinching frames are chosen for the 4-mer TR. (B) Overlapping repeats are defined as distinct TRs that only partially overlap each other. In contrast, fractalized repeats are defined as having a large *k* “parent” TR component and smaller *k* “fractal” TR components residing within each of the parent units. In this example of a hyper-fractalized repeat, a parent 11-mer repeat has repeat units containing smaller 5-mer repeats, each of which contain dinucleotide repeats. In addition to the hyper-fractalized repeat content, the 11-mer repeat contains an internal, non-compatible, non-fractal, 6-mer repeat, whose cinching would break homological integrity of columns representing the cinched 11-mer. Lowercase lettering (e.g., “*abab*”) highlights different repeat units within the 22 bp sequence. Below is the optimal, intermediate 2D self-alignment following *LINEUP* cinch-t and partial cinch-k (to *k*=5) cinching (see Fig. 3)

Distinct from overlapping repeats, there are fractalized repeats, which are “parent” repeats with internal repeats (see Fig. 2B for an example of a hyper-fractalized TR). Development of *LINEUP* revealed that TRs with divergent unit lengths produced by internal fractal repeats can be identified by considering direct tandem repeats that appear only in a sequence’s 2D consensus. This consensus of a 2D self-alignment approximates an ancestral, less-expanded sequence. This motivated the development of the *LINEUP* program as a series of cinching modules (Fig. 3). An initial “cinch-t” module cinches parent TRs of equal unit lengths while eschewing the cinching of the shorter, internal, repeats they may possess (Fig. 3A–3D). A subsequent “cinch-k” module cinches internal fractal TRs (Fig. 3C), which allows the identification of adjacent repeats of divergent lengths by a final “cinch-d” module (Fig. 3A–3C).

**Figure 3.**
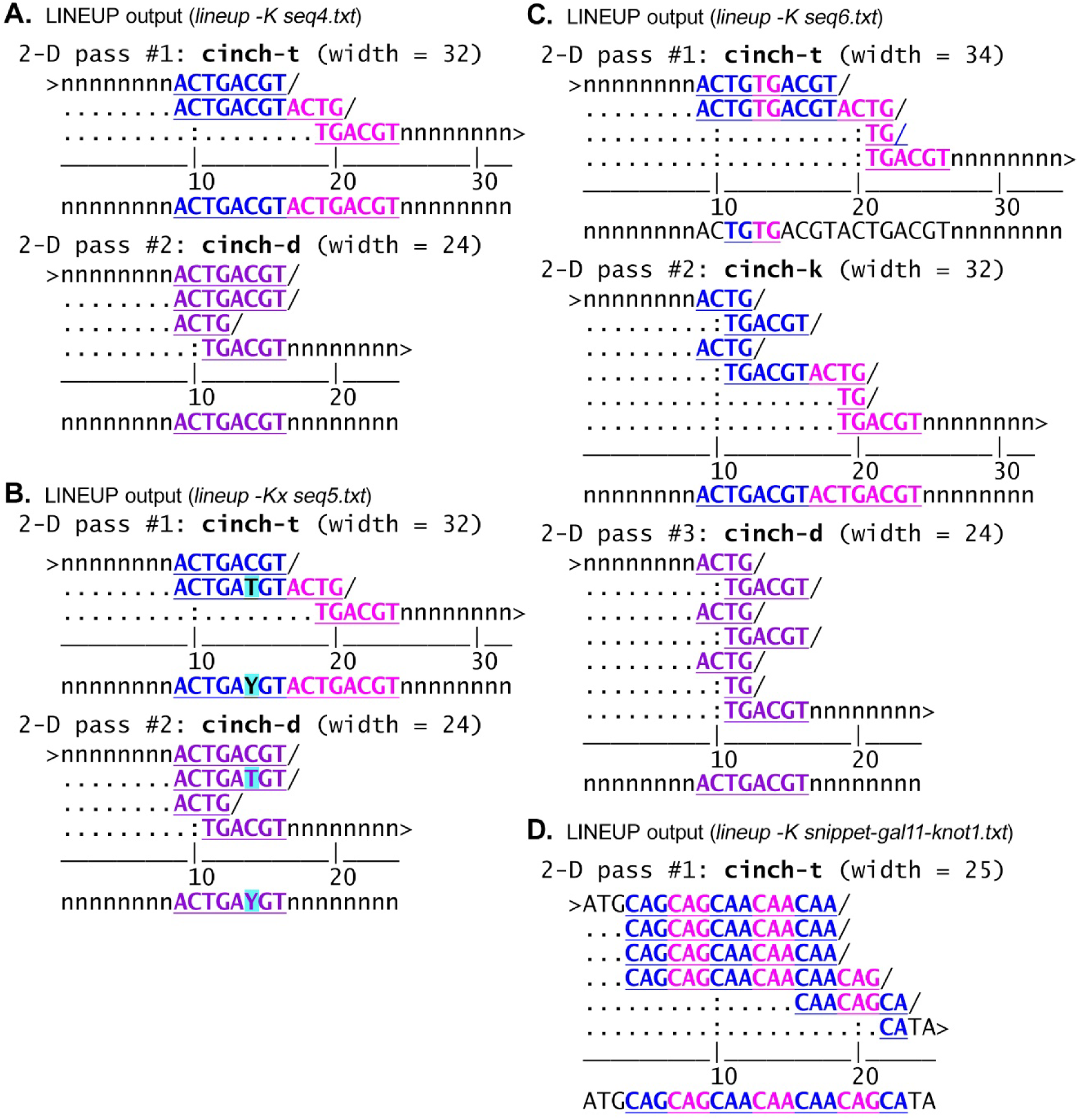
Cinching fractalized TR content of divergent unit lengths. (A) A sequence has three copies of an ancestral 8-mer sequence. Repeat unit three is not cinched by an initial “cinch-t” module of *LINEUP* due to a divergent length caused by an internal dinucleotide repeat (TG)_2_. The cinch-t module cinches up the two units of the 8-mer and separately cinches the dinucleotide repeat in the third 10 bp TR unit to produce an intermediate 2D alignment. The consensus row in this intermediate alignment reveals the presence of adjacent uncinched sequence homology. A subsequent “cinch-d” module cinches this divergent TR content to produce the final 2D alignment. (B) Shows 2D alignment steps for a sequence like (A) but differing by a single nucleotide transition (“Y”). (C) An intermediate “cinch-k” module to cinch internal k-mers, called fractal repeats of parent TRs, is required to make optimal use of cinch-d cinching in the consensus row after initial cinching of parent TRs by cinch-t. In this example, the units differ by having two or three copies of the dinucleotide fractal repeat. (D) The *LINEUP* program readily identifies the optimal 2D self-alignment of the nucleotide sequence encoding the polyglutamine-rich sequence MQ_23_HI in yeast *MED15* (*gal11*). Despite the presence of fractal repeats consisting of either (CAG)_2_ or (CAA)_3_, *LINEUP* first finds the most extensive parent repeat [(CAG)_2_(CAA)_3_]_4_ as well as the compatible overlapping repeat (CAACAG)_2_.

The *LINEUP* program can also (*i*) identify imperfect repeats differing by transitions (Fig. 3B), (*ii*) relax mononucleotide runs that did not aid cinching in the cinch-d module, (*iii*) confirm that the initial 1D starting sequence can be recovered from its 2D auto-alignment, and (*iv*) randomly shuffle input sequences before performing 2D self-alignment.

The capacity to randomly shuffle a sequence using the Fisher-Yates (FY) algorithm (Fisher and Yates 1953) is built into *LINEUP* and has two uses. First, FY shuffling was instrumental in identifying sequences that knotted into a pathological 2D alignment and led to further program development. This usage revealed that 2D micro-paralogical alignment is non-trivial in many ways. The second use of FY shuffling is for comparisons of biological sequences with their randomly shuffled versions, which possess the same mononucleotide base compositions. This second application is used extensively here.

It would be an oversimplification to describe *LINEUP* as a program for identifying TRs or even divergent TR content, although it can be used as such. This is the first example of a program designed to contract repeat-like content into a 2D self-alignment, which respects the homological basis of alignment columns. Ideally, if multiple nucleotides from the same sequence are placed into the same alignment column, it is because of an inference of homological equivalence. This type of program thus produces evolutionary-framed hypotheses of how repeat content has expanded and is related to each other. This is equivalent to the aims of phylogenetics in which MSAs are hypotheses of the homological relationships between nucleotide positions in related sequences. Second, *LINEUP* is the first program (albeit a sketch prototype) for pretreating sequences by 2D cinching before multiple sequence alignment. The closest comparable analog of this task might be repeat masking, but repeat masking is not concerned with 2D homological integrity, nor is it typically concerned with low copy number repeats. Last, the 2D cinching of TR content must be able to handle sequences characterized by repeat instability that typically evolve toward TR “knots” composed of multiple overlapping *k*-mers of different sizes. Figure 3D shows an example from a polyQ-encoding TR knot in yeast *MED15* (*gal11*), which is efficiently cinched by *LINEUP*.

Development of the *LINEUP* program followed four imperative goals. The first goal was to auto-align a large example corpus of biological sequences and artificial knotting sequences identified through millions of trials of random shuffling of biological and contrived sequences with extreme nucleotide compositions. The second goal was to minimize the average width cinch ratio (WCR) in example corpora, where WCR is defined as the final cinch-d 2D width divided by the initial sequence length. Thus, a WCR ratio for a sequence of 0.5 means the final 2D width was one-half the initial 1D sequence length. The third goal was to favor initial cinch-t cinching of TR content with equal unit lengths over cinching the same content later using the cinch-d module. The fourth goal was to favor a smaller and simpler code base than a more extensive code base (see Materials & Methods for algorithmic details). The current *LINEUP* program (v5.01) balances all four goals without compromising the 2D homological integrity of columns in any known sequence.

A fifth possible goal would be to achieve non-pathological cinching in the shortest computational time. This will require testing different programming approaches and is left as a future goal. Currently, the *LINEUP* program (v5.01) has cubic time complexity, *O*(*n*^3^), where *n* is the length of the input sequence (see Materials & Methods). However, the program is designed to be a 2D self-aligner of genomic block lengths typical of exonic CDS and non-protein-coding regulatory modules (e.g., *LINEUP* takes ∼0.5 s to process a 2 kb sequence on a laptop).

### A distinctive signature characterizes all protein-coding sequences

I now report how the *LINEUP* program reveals a fundamental signature of protein-coding sequence and its evolution. An initial goal was to use the *LINEUP* program to study a set of developmental enhancers in *Drosophila* that encode diverse responses to a developmental morphogen gradient. These enhancers are known as neurogenic ectoderm enhancers (NEEs) and share a complex TF binding site lexicon and regulatory grammar (Erives and Levine 2004). Interestingly, these enhancers evolve functional spacer elements that regulate the synergy between two transcriptional activators via the linkage between their binding sites (Crocker, et al. 2008). Furthermore, the divergence of functional spacers occurs via indels produced by TR instability (Crocker, et al. 2010; Brittain, et al. 2014). These enhancer sequences inspired the first 2D alignments, which were initially constructed by hand in exquisite detail and could be used as target benchmarks in developing computational 2D alignment. For this study, I use the first intron of the *ventral nervous system* defective (*vnd*) gene, which contains the oldest known NEE, as it is conserved in *Anopheles gambiae* (Erives and Levine 2004). I also use the *vnd* intronic sequence from 18 different *Drosophila* species, ranging in length from ∼1.5 to 3 kb (Table 1). These lengths roughly scale with their known respective genome sizes and represent an extreme case of evolutionary divergence via indels.

**Table 1.** Many protein-coding sequences are distinctively depleted in dinucleotide repeats and enriched in trinucleotide repeats.

| **Data set (n)** | **Range of lengths (bp)** | **cinch-t $k_2/k_3$ ratio** | **Sequence content** |
| --- | --- | --- | --- |
| *Drosophila vnd* CDS (18) | 917–980 | <u>1.04</u> | Encodes homeodomain and C-terminus |
| Endopterygota *gapdh* CDS (7) | 999–1,002 | <u>1.18</u> | Encodes NAD(P) binding and glyceraldehyde 3-phosphate dehydrogenase domains |
| Eukaryotic *MED15* CDS (4) | 2,250–3,963 | <u>1.30</u> | Encodes KIX domain and substantial low complexity peptide sequence, polyglutamine runs, and intrinsically disordered regions |
| *E. coli* K-12 *clpB* CDS (1) | 2,574 | <u>1.39</u> | Encodes two nucleotide-binding domains (NBDs); ATPase chaperone |
| Human *MYC* 20 kb intergenic (1) | 20,000 | 2.75 | Non-protein-coding intergenic sequence (hg38 chr8:127716001-127736000) |
| Randomly shuffled versions of *Dros. vnd* CDS (180 = 18 x 10) | 999 | 3.17 | Non-biological sequence from 10 Fisher-Yates reshufflings of *vnd* CDS 3 from 18 different *Drosophila* sp. |
| Randomly shuffled versions of *Dros. vnd* intron (180 = 18 x 10) | 999 | 3.28 | Non-biological sequence from 10 Fisher-Yates reshufflings of *vnd* intron 1 from 18 different *Drosophila* sp. |
| Randomly shuffled versions of *E. coli* *clpB* (10) | 2,574 | 3.40 | Non-biological sequence from 10 Fisher-Yates reshufflings of *E. coli* *clpB* CDS |
| *Drosophila vnd* intron (18) | 1,459–2,865 | 3.61 | ~500 bp neurogenic ectoderm enhancer (NEE) and fast-evolving non-protein coding/non-regulatory content |

Data sets are ordered by increasing $k_2/k_3$ TR content ratios, showing that the ratios for protein-coding sequences (CDS) constitute a distinctive lower-class range (underlined ratios).

To identify a suitable negative control data set for the *vnd* intronic data, I chose the CDS region from the terminal *vnd* exon from the same 18 *Drosophila* species. This “negative control” data set is thus: (*i*) located in the same genetic locus as the intronic region, (*ii*) endowed with a protein-coding reading frame unlike the regulatory and non-functional intronic sequences, and (*iii*) less divergent in length (917 to 980 bp, see Table 1). This region also encodes an NKX-class DNA binding homeodomain (60 aa) and C-terminus (total avg. length 308 aa).

Analysis of these two matched data sets (*vnd* intronic vs. protein-coding) from 18 different species of *Drosophila* reveals that their “cinch-able” repeat content, as measured by the *LINEUP* cinch-t module, is surprisingly quite comparable. Each data set averages a little over five cinched TRs per 100 bp (see Fig. 4A). This measurement, however, does not account for differences in repeat copy number. For example, much of the repeat content in the *vnd* protein-coding data comprises TRs with only two copies.

**Figure 4.**
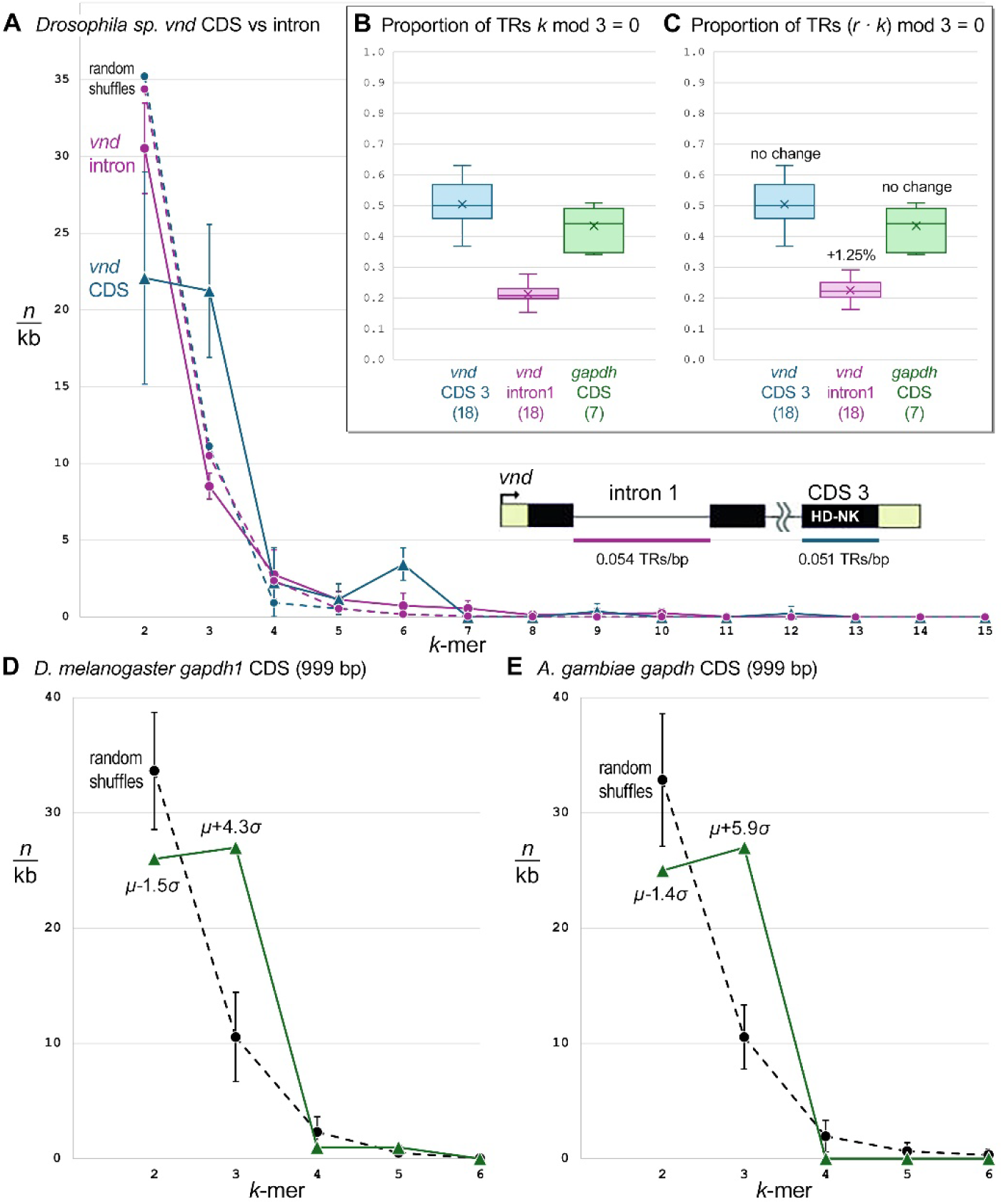
Protein-coding sequences are depleted in dinucleotide TRs and enriched in trinucleotide TRs. (A) TR content by *k*-mer size in two regions of the *vnd* locus from 18 *Drosophila* species is shown. One region is a protein-coding sequence (triangle data points, solid line) encoding an NKX-class homeodomain. The second is an intronic sequence (circle data points, solid line). Dotted lines represent *k*-mer content for 180 Fisher-Yates shuffled sequences (10x) from the *vnd* CDS (blue dotted line) and intronic (purple dotted line) data sets. (B, C) Protein-coding sequences are enriched in mod 3 TR content relative to non-coding sequences. Shown are the proportions of TR content with *k* mod 3 = 0 (B) or (*r* · *k*) mod 3 = 0 (C). The *vnd* sequences correspond to the CDS or intron sequence from 18 *Drosophila* species. The *gapdh* CDS sequences correspond to the ∼1 kb CDS sequence from 7 insects (5 flies, one moth, and one beetle). Despite *gapdh* not evolving by detectable indels at the amino acid level, it is enriched in *k* mod 3 TR content (B). Only the non-protein-coding sequences have additional TR content in which the product of the number of repeats *r* (not counting the first unit) times the *k*-mer size modulo 3 is zero (C). This suggests that the evolutionary path to three repeats (*i.e*., mod 3) for non-mod 3 *k*-mers in CDS regions sufficiently prohibits repeat expansion. (D, E) The *gapdh* coding sequence is enriched in trinucleotide tandem repeats and depleted in dinucleotide TRs (triangle data points). TR content by *k*-mer size is shown for the *gapdh* CDS of *D. melanogaster* (D, solid green line) or *A. gambiae* (E, solid green line). This TR content is contrasted with 25 trials of Fisher-Yates reshuffling of the corresponding sequence (circle data points on dotted line). Error bars denote +/− one standard deviation.

While the *vnd* homeodomain-encoding C-terminus had a surprising amount of elevated TR content, it nonetheless possessed a distinctive signature that distinguished it from the intronic region. The *vnd* protein-coding sequence is significantly depleted in dinucleotide repeats relative to trinucleotide repeats, even though its dinucleotide repeats still number slightly more than its trinucleotide repeats (Fig. 4A). This is to be expected because instability at dinucleotide-based TRs will produce indels disrupting the protein-coding reading frame and would represent mutation-prone alleles.

A dinucleotide sequence expanded to four copies (i.e., three extra copies) would constitute an insertion of 6 bp (3 x 2 base pairs), which would conserve a translational reading frame. To understand whether some of the dinucleotide repeats of the *vnd* CDS have a copy number that is compatible with a triplet reading frame, I analyzed the proportion of *k* mod 3 = 0 and (*r* · *k*) mod 3 = 0 content across the *vnd* protein-coding and non-coding sequences data sets, where *k* is the repeat unit length and *r* is the repeat number minus one (Fig. 4B–4C). Only the *vnd* intronic data set is seen to contain some (*r* · *k*) mod 3 = 0 content from non-mod 3 *k*-mers, suggesting that such content is eliminated and/or never generated in protein-coding sequence (Fig. 4C). This might be the case because three extra copies of any non-mod 3 *k*-mer could not be derived *de novo* without transitional repeat expansions of one to two copies, each of which would disrupt the codon frame.

Because the terminal *vnd* exon might allow an unusually high number of tandem repeats, perhaps downstream of the homeobox sequence, I then analyzed the *gapdh* CDS of various insects. The *gapdh* sequence was chosen because Gapdh is a highly conserved and highly expressed enzyme that does not evolve via indels that are apparent at the amino acid level. The *gapdh* CDS resides in a single exon in dipterans and lepidopterans. However, in a beetle outgroup sequence from *Anoplophora glabripennis*, chosen for its larger genome size of 980 Mb (McKenna, et al. 2016; McKenna 2018), it is interrupted by multiple introns. Because the protein sequence alignment of Gapdh for this clade is gapless, unlike the *vnd* protein-coding sequence, it was surprising to see that *gapdh* nucleotide sequences are still enriched in TR content characterized by *k* mod 3 = 0 (Fig. 4B–4C). However, much like the *vnd* protein-coding sequences were dinucleotide repeat-depleted (Fig. 4A), so were the *gapdh* protein-coding sequences (Fig. 4D–4E, solid lines).

To understand the origin of the unexpected repeat content and *k*-mer distribution for *gapdh*, I also compared it to Fisher-Yates random reshufflings of the CDS region from the two divergent lineages of *D. melanogaster* and *A. gambiae* (Fig. 4D–4E, dotted lines). This shows that trinucleotide repeats are enriched to levels 4 to 6 standard deviations above the mean of their shuffled counterparts (Fig. 4D–4E). In contrast, the dinucleotide repeats are depleted to levels more than one standard deviation below the mean of shuffled sequences. The enrichment of trinucleotide repeats relative to random sequences with identical nucleotide composition indicates that the trinucleotide repeats are biologically generated as opposed to a feature of random chance based on nucleotide frequencies. Furthermore, the depletion of dinucleotide repeats indicates extreme negative selection against non-mod 3 *k*-mers, whose instability would disrupt protein reading frames.

To determine whether the distinctive protein-coding signature left by RS during *gapdh* and *vnd* evolution is seen in genes encoding proteins with fast-evolving amino acid repeats, low complexity, and intrinsically disordered protein regions, I looked at the *MED15* gene. *MED15* encodes a subunit of the eukaryotic Mediator co-activator complex and is characterized by these protein features. I examined the *MED15* gene from two ecdysozoans (*D. melanogaster* and *C. elegans*) and two fungi (*S. cerevisiae* and *A. gossypii*). The length of this CDS from the four lineages ranges from ∼2.3 kb up to almost 4 kb (Table 1). I find that this *MED15* data set is also characterized by an extremely low dinucleotide to trinucleotide TR ratio of 1.30 (Table 1), similar to the ratios for the CDS data from *gapdh* (1.18) and *vnd* (1.04).

To determine whether the distinctive RS-generated protein-coding signature is an inherent feature of the amino acid codon system inherited from the latest universal common ancestor (LUCA) by all cellular life, I also looked at the CDS for the large *clpB* gene from the bacterium *E. coli*. The *clpB* gene is conserved across all domains of life, with an important exception being animals (Erives and Fassler 2015), and encodes an essential protein-folding chaperone with ATPase activity. This *clpB* gene is also characterized by dinucleotide repeat depletion and has a di- to tri-nucleotide TR ratio of 1.39 (Table 1). In contrast, randomly shuffled versions of *clpB* have a ratio of 3.40, similar to the ratios of shuffled *gapdh* CDS (3.15) and shuffled *vnd* intron (3.28), all of which are slightly below the *vnd* intronic ratio of 3.61 (Table 1).

To sample another biological (non-shuffled), non-protein-coding sequence, I measured the repeat ratio for a 20 kb region upstream of the human *MYC* locus, which has characteristically large intergenic regions. This region had a *k*_2_/*k*_3_ repeat ratio of 2.75, similar to that of the non-CDS data sets (Table 1). Thus, the five non-protein-coding data sets have a tight distribution around their average *k*_2_/*k*_3_ repeat ratio of 3.24, while the four protein-coding data sets have a similar tight distribution around their average ratio of 1.23 (Table 1).

### Trinucleotide repeats in protein-coding sequences do not favor the codon frame

To test the idea that the trinucleotide repeat enrichment in *gapdh* is a signature of the pervasive force of replication slippage acting even within a gene evolving in a “gapless” manner, I next investigated whether its trinucleotide repeats favor the codon reading frame. For example, the Gapdh enzyme-coding sequence typically has multiple sites of repeated amino acids. I first confirmed that different lineages have a substantial number of lineage-specific TRs, as would be the case if TR content is pervasively generated *de novo*. This was the case as 2/3 of the combined repeat content within the *D. melanogaster* and *A. gambiae gapdh* genes (or 1/3 each) is not shared (Table S1 and Fig. S1). This confirms that analyses of TR content in *gapdh* genes from divergent lineages are sampling mainly independent evolutionary RS events.

To determine if triplet repeats are more frequent in the codon frame, I looked at three divergent *gapdh* sequences from *Drosophila*, *Anopheles*, and the moth *Bombyx mori* (Table S2 and Fig. S2). Of all the trinucleotide repeats in these three lineages, 66% have a single cinching frame (46/70) and provide an unambiguous test data set for whether they coincide with the codon reading frame. Only ∼26.1% of these (12/46) coincide with the protein-coding frame (Table S2). This shows that the trinucleotide content is not enriched over the codon reading frame and conforms more closely to the null model (33.3%), whereby trinucleotide repeats are produced by replication slippage without regard to the protein-coding frame.

To determine the extent to which any repeated amino acids are associated with trinucleotide repeats, the nucleotide MSA was compared to the protein MSA of Gapdh (Figs. S2 and S3). This shows three essential things. First, at Gapdh alignment sites where an amino acid is repeated across many lineages (see Fig. S3), it is not the case that this is the result of conserved tandem repeats (see Fig. S2); TRs do not usually underlie such sites in all lineages, and even when they almost do so they are comprised of different repeat sequences. Second, at Gapdh sites where an amino acid is repeated only in a single lineage, it is almost always the case that a recently acquired TR is responsible. Third, many trinucleotide repeats are not associated with repeated amino acids (see underlined, non-repeating amino acids in Fig. S3). Collectively, these results establish that trinucleotide repeat enrichment in *gapdh* is RS-driven and occurs in all three reading frames.

### Fixation of RS-associated errors is pervasive even when protein-coding sequences appear to diverge without apparent indels

As shown, there is a universal signature for diverse protein-coding sequences in the TR content ratio of *k*_2_/*k*_3_ < 1.4, less than half the typical value for intergenic, regulatory, or randomly shuffled protein-coding sequences. Nonetheless, highly conserved genes such as *gapdh*, which evolve in a gapless manner, are still enriched in trinucleotide repeat content across all three triplet frames. Therefore, to determine how RS is persistently creating trinucleotide repeats in *gapdh* genes *de novo* without creating insertions detectable at the amino acid level, an inspection of the trinucleotide repeats in the MSA was undertaken. At many such sites, the 2D multiple sequence alignment provides a different and much simpler picture of mutational evolutionary events (Fig. S2 and Fig. 5).

**Figure 5.**
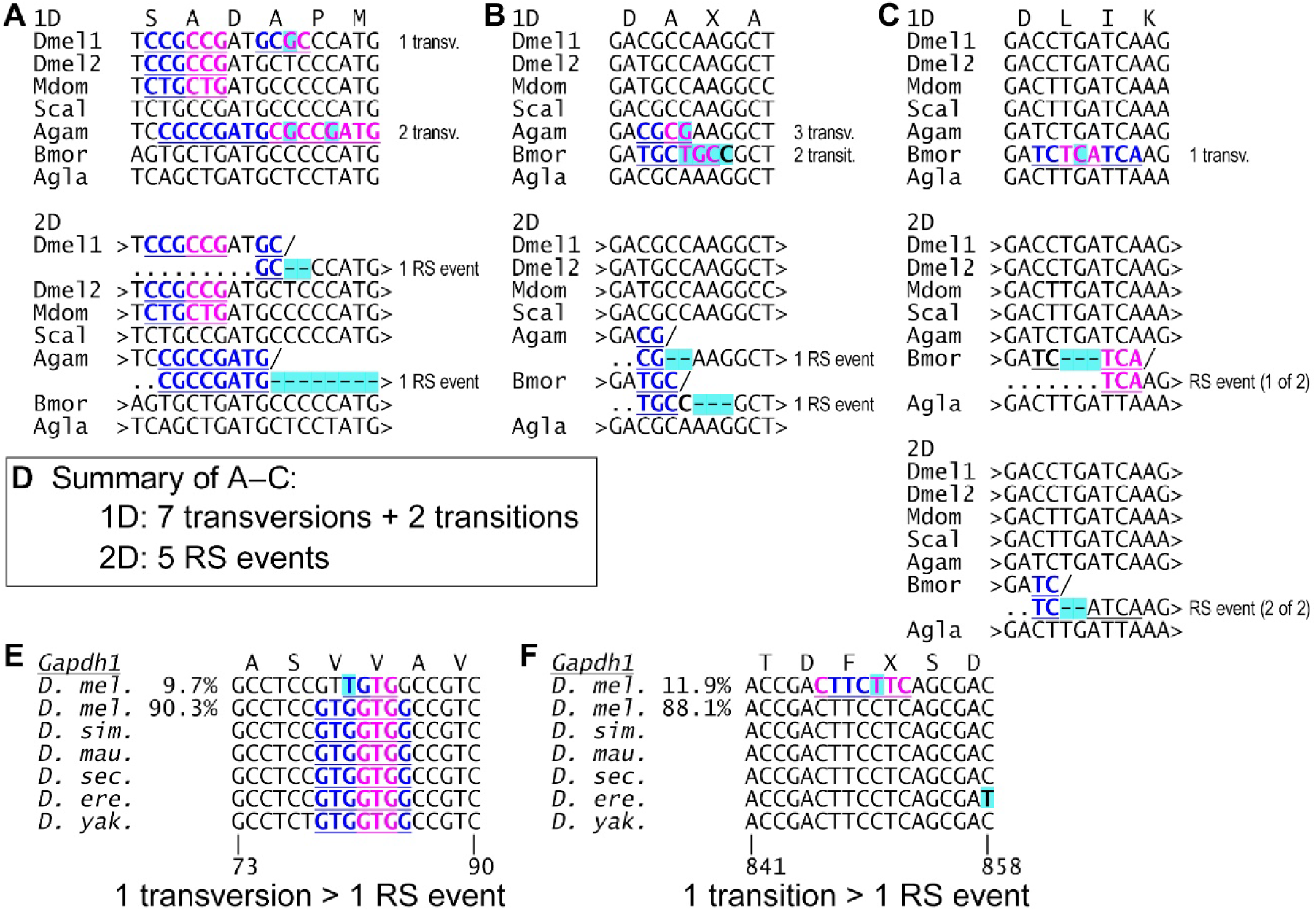
Cryptic replication slippage is responsible for most apparent transversions but also many transitions. (A–C) Shown are three example regions from the *gapdh* alignment in which 9 total point substitutions (7 transversions + 2 transitions, highlighted in cyan) are apparent in the 1D gapped alignment (top MSA in each panel). All nine substitutions transform into conserved ancestral nucleotides in their 2D alignments (bottom MSA in each panel), which infer five replication slip events based on tandem repeats in the regions of the highlighted substitutions. These replication slips are cryptic because selection only allows gapless RS events to persist in this highly conserved protein-coding sequence. These three examples demonstrate the power and versatility of modeling sequence evolution using both gaps and replication slips. In the third example in (C), two possible RS events can explain away the transversion seen in the 1D alignment. (D) Altogether, the 9 unambiguous TR-associated substitutions in the three examples from A–C transform into only five RS events. “Dmel1” and “Dmel2” = *Drosophila melanogaster Gapdh1* and *Gapdh2* genes, resulting from a melanogaster species group specific duplication (see Fig. S5); “Mdom” = *Musca domestica*; “Scal” = *Stomoxys calcitrans*; “Agam” = *Anopheles gambiae*; “Bmor” = *Bombyx mori*; and “Agla” = *Anoplophora glabripennis*. (E–F) Shown are two regions within the *Gapdh1* gene from the melanogaster species group. Both regions have SNPs from the Drosophila Genetics Reference Panel (DGRP). In each case, the major allele in *D. melanogaster* is ancestral and the minor allele is a derived change associated with the origination of a TR *de novo*. In E, the derived polymorphism is a silent transversion (G>T) associated with a dinucleotide repeat (TG)_2_. In F, the derived polymorphism is a non-synonymous transition (C>T) associated with a trinucleotide repeat (CTT)_2_C = C(TTC)_2_. Similar RS events pertain to 50% of all SNPs in the *Gapdh1* gene of *D. melanogaster* and include all apparent transversions (Fig. S6).

Three *gapdh* sites are representative of the mutational forces at play here and are chosen for exposition because they have only a few unambiguous lineage-specific substitutions (Fig. 5). In these three different regions spanning 4 to 6 codons each, the gapless 1D alignments show nine total unambiguous point substitutions that lie within perfect tandem repeats. Intriguingly, 78% of these (7/9) are transversions. However, in the 2D aligned versions, all nine substitutions become conserved ancestral residues of an adjacent tandem repeat. In short, the evolutionary selection of insertion-less Gapdh protein sequence has ordained that *de novo* tandem repeats persist only if they have replaced adjacent sequences. It is thus presumed that the immediate repair of *de novo* tandem repeats in the *gapdh* evolutionary regime frequently results in the deletion of either one copy of the new repeat or an adjacent sequence of similar length. However, only the latter RS-associated events would be observable.

To approximate a lower bound estimate on the proportion of apparent point substitutions that may originate with RS replacing adjacent sequence in *gapdh*, I first identified unambiguous, lineage-specific nucleotide substitutions. These are defined as mutations to a nucleotide that is not seen in any other *gapdh* sequence in the data set (7 divergent insect *gapdh* sequences). I then counted the subset of these unique substitutions occurring in *LINEUP*-identified TRs except the unpolarized differences occurring in the outgroup beetle sequence (Fig S2). Nearly 27% (57/214) of these unambiguous lineage-specific differences are embedded within perfect tandem repeats. This proportion is slightly more than expected given the repeat content (∼52 unique differences expected given the observed 24.2% repeat content), but this expectation ignores the role of negative selection in culling repeats.

To understand the lower-bound estimate on the percentage of RS-generated substitutions (27%) from a perspective of mutational mechanisms, I evaluated all 181 nucleotide differences between *D. melanogaster* and *A. gambiae gapdh* genes and calculated 50% for this data set (Fig. S4). To explain, half of these (90/181) lie within TRs and 2/3 of these TR-embedded differences (60/90) are transversion-type differences (Fig. S4). This two-fold excess of transversions over transitions is what would be expected by a null model of random chance because every nucleotide has two possible transversions but only one possible transition (Stoltzfus and McCandlish 2017). This further supports the conclusion that apparent point substitutions embedded in TRs are an artifact of misalignment in 1D gapped alignments.

The point differences within TRs in the *Drosophila*/*Anopheles* pair-wise comparison have a clear GC bias as C↔G (S) differences constitute the largest class at 46.7% (42/90), while A↔T (W) differences constitute the smallest class at 3.3% (3/90) (see bottom table in Fig. S4). In contrast, point differences outside of TRs have a smaller proportion of C↔G differences at 26.4% (24/91). Thus, the strong GC-bias, which is specific to TR-associated point differences, suggests that RS is assisted by the stronger bonding of C:G base pairs over A:T base pairs.

### One-half of all derived SNPs in D. melanogaster Gapdh1 are associated with new TRs and account for all transversion SNPs

The *gapdh* gene was duplicated in the melanogaster species group (see Fig. S5 for a phylogenetic tree of the *Gapdh1*/*Gapdh2* duplication), and so there are closely related species to *D. melanogaster* that can serve as outgroups for comparison. To look at population-level differences, I looked for regions within the *Gapdh1* gene having Drosophila Genetics Reference Panel (DGRP) SNP data (Mackay, et al. 2012; Huang, et al. 2014). There are 10 total SNPs genotyped in the isogenized DGRP lines at *Gapdh1* and the minor allele is the derived allele in 9/10 cases (see Fig. S6). Half of the SNPs (5/10) are C↔T (Y) transitions in which the derived allele is not associated with *de novo* TR origination. The derived alleles of the remaining 5/10 are all associated with *de novo* repeat creation. Considered as RS events, none of the TR-associated SNPs would count as substitutions. Considered as substitutions, they include two G↔T (K) transversions and three transitions (two Y and one R). Thus, half of the apparent point substitutions, including all the transversions, within *Gapdh1* of *D. melanogaster* may be the result of canonical base pairing with adjacent sequences during *de novo* TR generation by RS. Two of the five presumed RS-generated mutations are shown in Fig. 5E–F. One derived polymorphism is a synonymous transversion (G>T) associated with a dinucleotide repeat (TG)_2_ (Fig. 5E). The other derived polymorphism is a non-synonymous transition (C>T) associated with a trinucleotide repeat (CTT)_2_C = C(TTC)_2_ (Fig. 5F). The significance of these examples is that many population level differences are equivalently explained as duplications of adjacent sequence, the same as many (fixed) species-specific differences.

Altogether, these results suggest that replication slippage is a formidable force that changes the sequence of *gapdh* in a "gap-free" manner. A 2D alignment of *gapdh* shows that TR repeat content is created and maintained gap-free because new repeat units replace adjacent sequences (Fig. 5 and Fig. S2). In a 2D alignment, such sequences are restored to the same alignment columns, so they do not manifest in as many point substitutions. This includes many transversion mutations that are apparent only if the repeat copies are aligned to the replaced sequences.

### Using LINEUP to detect younger versus more mature TR content

The results reported here were identified with the *LINEUP* program, developed for 2D self-alignment of TR content. This program can also reveal aspects of TR evolution based on differential cinching by different *LINEUP* modules. Increased cinch-t cinching could indicate a greater proportion of young repeats (perfect repeats of uniform size), while increased cinch-d cinching could indicate a greater proportion of evolutionary mature repeat content that has experienced divergence in unit length through repeat fractalization. To determine whether repeat content in different data sets could be characterized as young (perfect TRs) versus old (more divergent TRs), I measured the cinching effect sizes for the serial *LINEUP* modules (cinch-t, cinch-k, and cinch-d) acting on the previously reported data sets.

Cinching effect sizes for the different *LINEUP* modules do bear some relation to specific types of data (Table 2). First, *gapdh* and randomized data sets have the lowest cinch-t effect sizes (blue shaded values in Table 2), consistent with high negative selection against non-mod 3 repeats in *gapdh* and an absence of RS-mediated TR creation in synthetic sequences. Secondly, and in contrast, protein-coding sequences that allow the evolutionary introduction of RS-associated indels have the highest cinch-t effect sizes, consistent with an abundance of young repeats (yellow shaded values in Table 2). Thirdly, the regulatory/non-protein-coding data sets had the greatest cinch-k effect sizes without having the greatest cinch-d effect sizes (red shaded values in Table 2). The former is due to extensive mononucleotide runs, while the latter might be because its repeat content is fractalized to an extent that is refractory to cinch-d. Nonetheless, when the cinch-k module is turned-off (“lineup -a0”) the *vnd* intronic data has a slightly higher cinch-d effect (−0.093) than the *vnd* CDS data (−0.086). This suggests there are aspects of cinching strategy that remain to be explored in future studies. Last, the final cinch-d WCR values were smallest for *MED15* and the non-protein-coding sequences, indicating the greatest cinching in sequences allowing the greatest amount of RS-generated TR content.

**Table 2.** Width cinch ratios indicate evolutionary mode.

| **Data set (n)** | **cinch-t**<br>**WCR** | **cinch-k**<br>**WCR** | **cinch-d**<br>**WCR** | **cinch-t**<br>**effect size** | **cinch-k**<br>**effect size** | **cinch-d**<br>**effect size** |
| --- | --- | --- | --- | --- | --- | --- |
| Endopterygota<br>*gapdh* CDS (7) | 0.884 | 0.637 | 0.496 | -0.116 | -0.247 | -0.141 |
| *Drosophila vnd* CDS<br>(18) | 0.833 | 0.634 | 0.475 | -0.167 | -0.199 | -0.159 |
| Eukaryotic *MED15*<br>CDS (4) | 0.838 | 0.594 | 0.436 | -0.162 | -0.244 | -0.158 |
| *Drosophila sp. vnd*<br>intron (18) | 0.868 | 0.602 | 0.455 | -0.132 | -0.266 | -0.147 |
| Human *MYC* 20 kb<br>intergenic (1) | 0.862 | 0.590 | 0.451 | -0.138 | -0.272 | -0.139 |
| Shuffled *gapdh* CDS<br>(7x10) | 0.886 | 0.645 | 0.506 | -0.114 | -0.241 | -0.139 |
| Shuffled *MED15*<br>CDS (4x10) | 0.876 | 0.628 | 0.480 | -0.124 | -0.248 | -0.148 |
| Shuffled *Dros. vnd*<br>CDS (18x10) #1 | 0.875 | 0.625 | 0.480 | -0.125 | -0.250 | -0.144 |
| Shuffled *Dros. vnd*<br>CDS (18x10) #2 | 0.872 | 0.624 | 0.479 | -0.128 | -0.248 | -0.145 |
| Shuffled *Dros. vnd*<br>intron (18x10) #1 | 0.880 | 0.639 | 0.498 | -0.120 | -0.241 | -0.140 |
| Shuffled *Dros. vnd*<br>intron (18x10) #2 | 0.880 | 0.641 | 0.502 | -0.120 | -0.239 | -0.139 |

## Discussion

Here, I introduced the concept of 2D representations to allow internal micro-paralogy to be represented in the same columns of a DNA sequence alignment. This is used to generalize GA into MPGA, which features a complete set of homology models (orthology + paralogy). In developing a prototype program called *LINEUP* to explore 2D sequence preparation, I found it helpful in generating hypotheses for how overlapping micro-paralogy is related. Its expanded multi-state homology model shows that 2D alignment is more powerful and versatile than the traditional 1D gapped alignment in which tandem repeats are left dispersed across different columns.

In using *LINEUP* to compare the TR content of non-protein-coding sequences to protein-coding sequences, I discovered an unexpectedly high amount of cryptic TR content in protein-coding sequences. This included genes, such as *gapdh*, that appear to evolve in an indel-free manner when analyzed with one-dimensional gapped alignment. Subsequent analyses revealed that RS produces abundant TR content in protein-coding sequence irrespective of the translational reading frame and that negative selection removes the non-mod 3 *k*-mer content. As this depletion is most noticeable for the most abundant *k*-mer bin of dinucleotide repeats, protein-coding sequences share a distinctively low ratio of 2-mers to 3-mers. This low ratio (∼1.2) is much lower than either non-protein coding sequences or randomly shuffled biological sequences (∼3.2).

The new perspectives provided by 2D alignment are profound and indicate that future studies are needed. Here, I showed how many apparent transversion mutations and runs of multiple point substitutions transform into conserved, ancestral nucleotide bases when sequences are aligned by 2D alignment. Thus, traditional GA has likely caused an inflated estimate of point substitution rates at the expense of indel rates. Furthermore, the transition to transversion ratio is likely much higher than estimated. This suggests that “in place” base pairing errors might pertain predominantly to transition mutations between chemically similar bases (i.e., between purines A↔G or between pyrimidines C↔T).

As shown here, 2D alignment can improve measures of sequence divergence and its classification (e.g., substitutions vs. indels, or transitions vs. transversions). However, another important result may be the enrichment of trinucleotide repeats in protein-coding sequences relative to randomly shuffled sequences sampled from the same sequences (Fig. 4). This trinucleotide enrichment occurs irrespective of the codon reading frame and regardless of whether a sequence encodes single amino acid repeats or not. I previously suggested that this is indicative of the pervasive force of replication slippage at the very least. However, this is not entirely correct because this point can be derived directly from the *k*_2_/*k*_3_ ratio of CDS data. Furthermore, were the trinucleotide enrichment merely a result of RS effects that escaped negative selection (unlike the dinucleotide repeat content), we would expect a similar enrichment in the *vnd* intronic data relative to their shuffled counterparts, and this is not observed (Fig. 4). This suggests, perhaps, that the evolutionary maintenance of protein-coding sequences may be assisted by the positive selection of alleles produced by replication slippage itself, particularly if transversions would be required to produce a beneficial allele. This would add to the list of examples in which mutational biases influence adaptive evolutionary processes (Cano, et al. 2023).

The challenges of computing 2D self-alignments are like those encountered by DNA replication and repair enzymes. Computationally aligning a complex TR knot is challenging for the same reason such sequences have a propensity for biological instability. Both challenges stem from the potential for base pairing with adjacent sequences. Therefore, it can be argued that base pair complementarity, the property underlying replication and transcription, is inherently mutagenic. Thus, it will be interesting to determine the correlation between TR biology and mutational hotspots in diverse sequences using 2D alignment methods.

## Materials and Methods

### *LINEUP* program development

*LINEUP* was written in C with an extensive set of command line options and is compatible with Linux and macOS systems. The *LINEUP* code, demo and test scripts, and an extensive example corpus are available on the GitHub code repository at https://github.com/microfoam/lineup. The *LINEUP* cinch-t module uses a dynamic programming method analogous to those used in global and local pairwise alignment (Needleman and Wunsch 1970; Smith and Waterman 1981). The *LINEUP* method is to traverse through a grid, which scores a sequence to itself and delineates tandem repeats for cinching. *LINEUP* also uses C structures to record compatible cinches in a 1D data format, which is used to guide initial cinching. The *LINEUP* cinch-k, cinch-d, and other related modules work on two-dimensional array representations.

### LINEUP’s cinching corpus

Although there are no outstanding or unsolved cinching problems, *LINEUP* is built to write to a file with details of problem sequences it encounters (“waves/foam_and_chowder.log”). In early program development, cinching problems were of two major types: (*i*) 2D self-alignments in which non-paralogous nucleotides were placed into the same column (“bad slips”), and (*ii*) 2D self-alignments in which the original 1D sequence could not be perfectly recovered from the final 2D alignment. Originally, cinching problems were identified from biological sequences. Later, *LINEUP* scripts to engage Fisher-Yates shuffling of experimental sequences were used to generate and 2D self-align tens of millions of synthetic sequences 100 bp in length over the lifetime of program development (e.g., the “surfboard-washingmachine” script). The generation of short sequences facilitated the identification of problem cinching knots to within 100 kb. This led to the discovery of hyper-fractalized repeats, overlapping repeats, bad slips, cycling knots, artifactually large *k*-mer size repeats that should be cinched at smaller *k*-mer sizes, and numerous other issues.

Near the end of program development, problematic sequences were identified at a rate of 1 in a million 100 bp DNA sequences produced by random sampling of extreme nucleotide compositions and 2D self-aligned using extreme *LINEUP* run parameters (e.g., “lineup -x”, which lowers the threshold for transition matching). This was used to build the extensive example corpus of cinching problems that *LINEUP* was designed to solve (see the “wave” sub-directories “snippets/”, “squeegee/”, “tubespit/”, and “chowder/” in the repository). The entire example corpus can be run from test scripts sharing the base name "surfboard-cleanup_set-".

### LINEUP time complexity

To determine the computational time complexity of the *LINEUP* program a script was written to generate 2 kb, 4 kb, 6 kb, 8 kb, and 10 kb randomly shuffled strings from each of the *vnd* CDS and *vnd* intron from five divergent *Drosophila* species (*D. melanogaster*, *D. pseudoobscura*, *D. willistoni*, *D. virilis*, and *D. grimshawi*). Thus, each size class consisted of 10 different sequences. The compute time for each class included Fisher-Yates shuffling, which has a time complexity ≤ O(*n*^2^). Considering the first data set of 20 kb (10 x 2 kb) as 1x input size taking 1x compute time, the five different time classes (*n* = 1–5) consistently had run times of *n*^3^ (1^3^), *n*^3^ − *n*/2, *n*^3^ − *n*, *n*^3^ − *n*, *n*^3^ + *n*^2^ − *n*, respectively, hence *O*(*n*^3^) or cubic time complexity.

### Measuring repeat content

All repeat content analyses in this study used *LINEUP* version 5.01 and relevant command line options. The examples in Figures 1–3 can be run from the script “surfboard-demo_toys” in the program’s GitHub repository.

## Acknowledgments

I thank John Reinitz and John Logsdon for helpful feedback on this project.

## Notes

https://github.com/microfoam/lineup

### Competing Interest Statement

The authors have declared no competing interest.

### Summary of Updates

The initial bioRxiv preprint was updated to match the version currently under review at a peer-reviewed journal. The title, abstract, and formatting have been updated as well as a few places in the text. The results, figures, tables, and associated code are the same as the original submission to bioRxiv.

