## Supporting Information (suplementary figures and tables) for "2D representations of DNA sequence show that most transversions are misaligned nucleotides associated with replication slippage"

**Figure S1. Most tandem repeats in *gapdh* are lineage specific.** Yellow highlighted TRs correspond to unique lineage-specific TRs in this pairwise comparison: 27/54 TRs from *D. melanogaster* and 27/54 TR from *A. gambiae*. 27/81 total number of distinct TRs are shared. Thus, roughly 1/3 is unique to each species (2/3 total), and 1/3 is shared.

```
4:      ....................................................babab...  1
        .....................................................abcabca  2
Dmel    ATGTCGAAGATCGGAATTAACGGATTTGGCCGCATCGGCCGCTTGGTGCTCCGCGCCGCC
Agam    ATGTCGAAGATCGGAATTAACGGATTTGGCCGCATCGGTCGTCTGGTGCTGCGCGCCGCC
        .....................................abcabc...abcabc........  1,2
        ..................................................ababab....  3
        .....................................................abcabca  4

3:      ..........abab....abcabca...................................  3,4
Dmel    ATCGATAAGGGCGCCTCCGTGGTGGCCGTCAACGATCCCTTCATCGATGTCAACTACATG
Agam    ATCACCAAGGGTGCGTCGGTGGTCGCCATCAACGATCCGTTCATCGGCGTCGACTACATG
        .................abcabc.....................................  5

3:      ................abcdeabcdeab................................  5
Dmel    GTTTACCTGTTTAAATTCGACTCGACTCACGGTCGTTTCAAGGGCACCGTTGCGGCTGAG
Agam    GTGTACCTGTTCAAGTACGACTCGACGCACGGTCGCTTCAAGGGCGAGGTGTCCGCCCAG
        abab............................................abab........  6,7

5:      ..........abcabc.........abcabc.....abab..abcdabcd.............  6,7,8,9
        ...............................................ababa...........  10
Dmel    GGCGGATTCCTGGTGGTGAACGGCCAGAAGATCACCGTGTTCAGCGAGCGCGACCCGGCCAAC
Agam    GACGGCTGCCTGGTCGTGAACGGCCAGAAGATTGCCGTGTTCCAGGAGCGCGACCCGAAGGCG
        .........................abcabc.....abab.......ababa...........  8,9,10

5:      ..............................bcdbcdb...........abab.abcabcab..  11,12,13
Dmel    ATCAACTGGGCCAGTGCTGGAGCCGAGTATGTGGTGGAGTCCACCGGAGTGTTCACCACCATT
Agam    ATCCCGTGGGGTAAGGCCGGCGCGGAGTACGTCGTTGAGTCGACCGGTGTGTTCACCACGACC
        ...................ababa.....abcabc...........ababab.abcabc....  11,12,13,14

6:      ........................abcabca..abcabc.abcabcab.........  14,16,17
        ............................abab..............abab.......  15,18
Dmel    GACAAGGCGTCCACCCACTTGAAGGGCGGCGCCAAGAAGGTCATCATCTCGGCCCCA
Agam    GAGAAGGCATCCGCCCATCTGGAGGGTGGCGCCAAGAAGGTCATCATCTCGGCCCCG
        abab........................abab.abcabc.abcabcab.........  15,16,17,18
        ..............................................abab.......  19

7:      .abcabc..abab........ababa..................................  19,20,21
        .........................abcabc.............................  22
Dmel    TCCGCCGATGCGCCCATGTTCGTGTGCGGCGTTAACCTGGACGCCTACAGCCCCGACATG
Agam    TCCGCCGATGCGCCGATGTTCGTCGTCGGTGTCAACCTGGAGGCGTACGAGCCATCGATG
        .abcabc..abab......abcabcabc................................  20,22,23
        ..abcdefghabcdefgh..........abab............................  21,24

6:      ..abcabc..............................................  23
        .......bcbc............abcabcab.......................  24,25
Dmel    AAGGTGGTCTCCAACGCCTCGTGCACCACCAACTGCCTGGCTCCCCTGGCCAAG
Agam    AAGGTCGTGTCGAACGCGTCCTGCACCACCAACTGTCTCGCCCCGCTGGCCAAG
        ......abab....abab.....abcabcab....abab...............  25,26,27,28
```

**Fig. S1 cont.** (part 2 of 2)

```
9:      .abcabc...........abab............abcabc.................... 26,27,29
        ......................abcabc...........cdecde.......abcabca. 28,30,31
Dmel    GTCATCAATGACAACTTCGAGATCGTCGAGGGTCTGATGACCACCGTGCACGCCACCACT
Agam    GTCATCAACGACAACTTCGGCATCCTGGAGGGTCTGATGACGACGGTGCACGCGACCACC
        .abcabc...........................abcabc..........abab...... 29,30,32
        ......................................bcdbcd..........abcabc 31,33

5:      ..........................abab.......abab...................... 32,33
        .........................................ababa..........abcabca 34,35
Dmel    GCCACCCAGAAGACCGTCGACGGTCCCTCTGGCAAACTGTGGCGCGATGGACGTGGCGCCGCC
Agam    GCCACCCAGAAGACGGTCGATGGCCCCTCGGGCAAGCTGTGGCGTGATGGCCGTGGTGCCGCC
        .....................................abab...........abcabc..... 34,35
        .........................................................bcabca 36

5:      .....abcabca...cadcad......abcabc.....abab..................... 36,37,38,39
Dmel    CAGAACATCATCCCGGCCGCCACCGGAGCCGCCAAGGCTGTGGGCAAGGTCATCCCCGCCCTG
Agam    CAGAACATCATTCCGGCGGCGACCGGTGCCGCCAAGGCGGTGGGCAAGGTCATCCCGGCCCTG
        .....abcabc..abcabcab......abcabc.............................. 37,38,39

4:      ...........................abab...............abab...abab... 40,41,42
Dmel    AACGGCAAGCTGACCGGCATGGCTTTCCGCGTGCCCACGCCCAATGTCTCCGTTGTGGAT
Agam    AACGGCAAGCTGACCGGTATGGCGTTCCGCGTCCCGACCCCGAACGTCTCGGTCGTCGAT
        ...........................abab...............abab.abcabca.. 40,41,42

1:Dmel    CTTACCGTCCGCTTGGGCAAGGGAGCCACCTATGACGAAATCAAGGCTAAGGTC
Agam    CTGACCGTGCGCCTGTCCAAGCCGGCCACCTACGACCAGATCAAGCAGAAGGTG
        ........abab.......................................... 43

7:      abcabca.................................abab....abcabca...cdcd. 43,44,45,47
        .....................................................abcabc.... 46
Dmel    GAGGAGGCCTCCAAGGGACCCCTGAAGGGAATCCTGGGCTACACCGATGAGGAGGTGGTCTCC
Agam    AAGGAGGCCGCCAACGGGCCGATGAAGGGCATCCTGGACTACACCGAGGAGGAGGTCGTGTCG
        .abcabc.................................abab.abcabcabca..bcbc.. 44,46,47,48
        ......cdecde................................................... 45

5:      ......abcabc....abab.........ababa.......................... 48,49,50
Dmel    ACCGACTTCTTCAGCGACACCCATTCGTCTGTGTTCGACGCCAAGGCTGGCATTTCGCTG
Agam    ACCGACTTCGTCGGCGACTGCCACTCGTCCATCTTTGACGCGAAGGCTGGCATCCAGCTG
        .......bcdbcd.........................abab.................. 49,50

2:      ......................abab......................... 51
Dmel    AACGATAAGTTCGTCAAGCTAATCTCGTGGTACGACAACGAGTTCGGTTAC
Agam    AGCGACACGTTCGTCAAGCTGATCTCCTGGTACGACAACGAGTACGGCTAC
        ....abab..............abab......................... 51,52

4:      ......abab.................ababa..abab.......... 52,53,54
Dmel    TCCAACCGCGTCATCGACCTGATCAAGTATATGCAGAGCAAGGACTAA
Agam    TCGAACCGCGTCGTCGATCTGATCAAGTACATGCAGACCAAGGATTAA
        ......abab...................................... 53
        ........abcabc.................................. 54
```

81 total

**Figure S2. About 27% of all lineage-specific point substitutions within the *gapdh* coding sequence are found within perfect tandem repeats.** Many more are found within imperfect tandem repeats differing from adjacent units by transition substitutions. Additional tandem repeats can be traced to have occurred in earlier ancestors. Another point to consider is that there is a strong signature of selection against dinucleotide-based TRs (Fig. 4A, 4D–4E). Altogether, these results point to a substantial proportion of point mutations being caused by RS-mediated, intralocus, micro-paralogical gene conversion. In a real sense, DNA sequence is inherently mutagenic. The shortest nucleotide sequences (< 10 bp) tend to copy themselves into adjacent sequences. This likely occurs when nascent tandem repeats are immediately repaired with the excision of adjacent sequences.

```
        M  x  K  I  G  I  N  G  F  G  R  I  G  R  L  V  L  R  A  x
Dmel1 >ATGTCGAAGATCGGAATTAACGGATTTGGCCGCATCGGCCGCTTGGTGCTCCGCGCCGCC>  0
Dmel2 >ATGTCGAAGATTGGTATCAATGGATTTGGTCGCATCGGCCGCTTGGTTCTCCGCGCCGCC>  0
Mdom  >ATGTCCAAAATCGGTATTAATGGATTCGGTCGTATTGGCCGTTTGGTTTTGCGTGCCGCC>  1
Scal  >ATGTCCAAAATCGGTATTAATGGATTCGGTCGCATTGGTCGCCTCGTCTTGAGGGCTGCC>  3
Agam  >ATGTCGAAGATCGGAATTAACGGATTTGGCCGCATCGGTCGTCTGGTGCTGCGCGCCGCC>  0
Bmor  >ATGTCAAAAATTGGAATCAATGGATTTGGCCGCATTGGCCGTTTGGTGCTCCGTGCTTCT>  1
Agla  >ATGGTGAAAATTGGAATCAACGGTTTTGGAAGAATTGGACGTTTGGTCCTCCGAGCATCT>  8
       ..|..|..|..|..|..|..|..|..|..|..|..|..|..|..|..|..|..|..|..|

        I  x  x  G  A  x  V  V  A  x  N  D  P  F  x  x  x  x  Y  M
Dmel1 >ATCGATAAGGGCGCCTCCGTGGTGGCCGTCAACGATCCCTTCATCGATGTCAACTACATG>  2
Dmel2 >ATTGATAAGGGCGCCAACGTTGTGGCCGTCAACGATCCCTTCATCGATGTGAACTACATG>  1
Mdom  >ATCGATAAGGGCGCCCAGGTTGTAGCCGTCAACGATCCCTTCATCGATGTCAACTACATG>  0
Scal  >ATTGATAAGGGCGCCCAAGTTGTGGCTGTCAACGATCCCTTCATCGATGTGGAATACATG>  1
Agam  >ATC/
       ACC---AAGGGTGC/
       ............GTC--GGTGGTCGCCATCAACGATCCGTTCATCGGCGTCGACTACATG>  5
Bmor  >ATTGAAAAGGGAGCTCAAGTGGTCGCTATAAATGACCCTTTCATCGGTCTTGACTATATG>  4
Agla  >ATTGAACGTGGAGCAGAAGTAGTAGCTGTAAATGATCCCTTCCTTGATGTTGATTATATG>  9
       ..|..|..|..|..|..|..|..|..|..|..|..|..|..|..|..|..|..|..|..|

        V  Y  L  F  K  x  D  S  T  H  G  R  x  K  G  x  V  x  x  x
Dmel1 >GTTTACCTGTTTAAATTCGACTCGACTCACGGTCGTTTCAAGGGCACCGTTGCGGCTGAG>  1
Dmel2 >GTCTACCTGTTCAAGTTCGATTCGACCCACGGACGTTTTAAGGGCACCGTTGCCGCCGAG>  0
Mdom  >GTCTACTTGTTCAAATTCGACTCCACCCATGGTCGTTTCAAGGGTACCGTTGCCGCTGAA>  0
Scal  >GTTTACTTGTTTAAATTCGATTCGACCCACGGTCGTTTCAAGGGTACCGTTGCTGCCGAA>  0
Agam  >GTGTACCTGTTCAAGTACGACTCGACGCACGGTCGCTTCAAGGGCGAGGTGTCCGCCCAG>  5
Bmor  >GTTTATCTTTTCAAGTATGATTCCACCCATGGCCGTTTTAAGGGCAGTGTTGAGGTTCAG>  4
Agla  >GTGTATCTCTTCAAATTTGATTCAACCCATGGACGTTACAAGGGTTGTGTGTCATCTGAT>  4
       ..|..|..|..|..|..|..|..|..|..|..|..|..|..|..|..|..|..|..|..|

        x  x  x     L  V  V  x  G  x  x  I  x  V  F  x  E  R  D  P  x   x
Dmel1 >GGCGG---ATTCCTGGTGGTGAACGGCCAGAAGATCACCGTGTTCAGCGAGCGCGACCCGGCC AAC>  0
Dmel2 >GGCGGT---TTCCTGGTGGTCAACGGCCAGAAGATCACCGTCTTCAGCGAACGCGACCCGGCC AAC>  1
Mdom  >GGGGGTAAAT---TGGTTGTGAACGGTCAGGCCATCACCGTTTTCAGTGAACGCGACCCCGCC AAC>  0
Scal  >GGTGGCAAAT---TGGTTGTTAACGGTCAAGCCATCACCGTTTTCAGCGAACGCGACCCCGCT AAC>  1
Agam  >GACGGC---TGCCTGGTCGTGAACGGCCAGAAGATTGCCGTGTTC/
       ...........................................CAG-GAGCGCGACCCGAAG GCG>  2
Bmor  >GATGG---ATTCCTTGTTGTTAACGGTAACAAAATTGCCGTTTTC/
       ..........................................TCA--GAAAGGGACCCTAAG GCC>  4
Agla  >GGAAAGAACT---TGGTAGTCGATGGCAAAAAAATTGCTGTATTC/         intron 1
       ...........................................CAA-GAAAGAGATCCCAAG^GCC> 12
       ..|..|..|..|..|..|..|..|..|..|..|..|..|..|..|..|..|..|..|..|  ..|
```

**Fig. S2 cont.** (2 of 4 pages)

```
          I  x  W  x  x  x  G  A  x  Y  x  V  E  S  T  G  V  F  T  T  x
Dmel1  >ATCAACTGGGCCAGTGCTGGAGCCGAGTATGTGGTGGAGTCCACCGGAGTGTTCACCACCATT>  2
Dmel2  >ATCAACTGGGCCAGCGCTGGTGCCGAATACATCGTGGAGTCCACTGGCGTGTTCACCACCATC>  1
Mdom   >ATCAACTGGGCCAGCGCTGGTGCCGAATATGTTGTTGAATCCACTGGTGTCTTCACCACCATC>  0
Scal   >ATTAATTGGGCCAGCGCTGGAGCCGAATATGTTGTGGAATCCACTGGCGTCTTCACCACCATC>  0
Agam   >ATCCCGTGGGGTAAGGCCGGCGCGGAGTACGTCGTTGAGTCGACCGGTGTGTTCACCACGACC>  7
Bmor   >ATTCCATGGGGAAAAGCTGGGGCTGAATATGTTGTAGAGTCTACTGGTGTCTTTACCACTACA>  7
Agla   >ATTCCATGGG/
         .........GACAGT-CGGGCGCTGATTACGTTGTTGAATCTACCGGAGTATTCACAACCATT>  3
         ..|..|..|..|..|..|..|..|..|..|..|..|..|..|..|..|..|..|..|..|..|

          x  K  A  S  x  H  x  x  G  G  A  K  K  V  I  I  S  A  P
Dmel1  >GACAAGGCGTCCACCCACTTGAAGGGCGGCGCCAAGAAGGTCATCATCTCGGCCCCA>  1
Dmel2  >GACAAGGCATCCACTCACTTGAAGGGCGGTGCCAAGAAGGTTATCATCTCGGCCCCA>  0
Mdom   >GACAAGGCTTCTGCCCATTTCAAGGGTGGTGCCAAGAAGGTCATCATCTCCGCTCCA>  1
Scal   >GACAAGGCCTCCGCCCATTTCAAGGGTGGTGCCAAGAAGGTCATCATCTCTGCTCCA>  1
Agam   >GAGAAGGCATCCGCCCATCTGGAGGGTGGCGCCAAGAAGGTCATCATCTCGGCCCCG>  3
Bmor   >GATAAAGCATCTGCTCACTTGGAGGGAGGTGCTAAAAAAGTTATTATATCAGCTCCC> 10
Agla   >GAAAAGGCTTCTGCTCACTTGGCTGGTGGAGCAAAGAAGGTTATCATCTCTGCTCCT>  6
         ..|..|..|..|..|..|..|..|..|..|..|..|..|..|..|..|..|..|..|

          S  A  D  A  P  M  x  V  x  G  V  N  L  x  A  Y  x  P  x  x
Dmel1  >TCCGCCGATGCGCCCATGTTCGTGTGCGGCGTTAACCTGGACGCCTACA-GC/
         ...................................................CCCGACATG>  1
Dmel2  >TCCGCCGATGCTCCCATGTTCGTTTGCGGCGTCAACTTGGATGCCTACAAGCCCGACATG>  0
Mdom   >TCTGCTGATGCCCCCATGTTCGTGTGCGGTGTCAACTTGGATGCCTACAAGCCAGACATG>  0
Scal   >TCTGCCGATGCCCCCATGTTTGTATGCGGTGTCAACTTGGATGCCTACAAACCAGACATG>  2
Agam   >TCCGCCGATGCGCCGATGTTCGTCGTCGGTGTCAACCTGGAGGCGTACGAGCCATCGATG>  4
Bmor   >AGTGCTGATGCCCCCATGTTTGTTGTGGGTGTTAACCTAGAAGCTTATGACCCCTCTTTT>  8
Agla   >TCAGCTGATGCTCCTATGTATGTCTGTGGTGTTAACCTTGATGCTTACAACCCAGCTGAC>  6
         ..|..|..|..|..|..|..|..|..|..|..|..|..|..|..|..|..|..|..|..|

          K     V  x  S  N  A  S  C  T  T  N  C  L  A  P  L  A  K
Dmel1  >AAG_GTGGTCTCCAACGCCTCGTGCACCACCAACTGCCTGGCTCCCCTGGCCAAG>  0
Dmel2  >AAG_GTGGTCTCCAACGCATCGTGCACCACCAACTGCTTGGCTCCTCTGGCCAAG>  2
Mdom   >AAA GTCGTTTCCAACGCTTCCTGCACCACCAACTGCTTGGCTCCCTTGGCCAAG>  0
Scal   >AAA GTGGTCTCCAACGCCTCCTGCACCACCAACTGCTTGGCTCCCTTGGCCAAG>  0
Agam   >AAG GTCGTGTCGAACGCGTCCTGCACCACCAACTGTCTCGCCCCGCTGGCCAAG>  4
Bmor   >AAG GTCATCTCAAATGCTTCTTGCACCACAAACTGTCTTGCCCCACTTGCAAAG>  5
Agla   >AAG^GTCATTTCAAATGCTTCATGCACAACAAATTGTTTGGCTCCCTTGGCCAAG>  3
         ..|  ..|..|..|..|..|..|..|..|..|..|..|..|..|..|..|..|..|

          V  I  x  D  N  x  x  I  x  E  G  L  M  T  T  V  H  A  T  T
Dmel1  >GTCATCAATGACAACTTCGAGATCGTCGAGGGTCTGATGACCACCGTGCACGCCACCACT>  0
Dmel2  >GTGATCAACGACAACTTCGAGATCGTCGAGGGTCTGATGACCACCGTTCATGCCACCACC>  1
Mdom   >GTTATCCACGATAACTTCGAAATCGTTGAAGGTCTCATGACCACTGTCCATGCCACCACT>  2
Scal   >GTTATCCACGACAACTGGGAAATTGTTGAGGGTCTCATGACCACTGTGCATGCCACTACT>  3
Agam   >GTCATCAACGACAACTTCGGCATCCTGGAGGGTCTGATGACGACGGTGCACGCGACCACC>  7
Bmor   >GTTATTCATGATAACTTTGAAATTGTTGAGGGCTTGATGACTACTGTTCATGCCACAACT>  5
Agla   >GTCATCCATGACAACTTTGAGATCGTTGAGGGTCTTATGACCACAGTACATGCCACCACT>  3
         ..|..|..|..|..|..|..|..|..|..|..|..|..|..|..|..|..|..|..|..|
```

**Fig. S2 cont.** (part 3 of 4)

```
         A  T  Q  K  T  V  D  G  P  S  G  K  L  W  R  D  G  R  G  A  x
Dmel1  >GCCACCCAGAAGACCGTCGACGGTCCCTCTGGCAAACTGTGGCGCGATGGACGTGGCGCCGCC>  1
Dmel2  >GCTACCCAGAAGACCGTCGATGGACCTTCCGGCAAGTTGTGGCGTGATGGACGTGGCGCTGCC>  0
Mdom   >GCCACCCAAAAGACCGTTGACGGTCCCTCCGGCAAATTGTGGCGTGATGGCCGTGGAGCTGCC>  1
Scal   >GCCACACAAAAGACCGTTGACGGACCCTCTGGCAAATTGTGGCGTGATGGTCGTGGTGCTGCC>  0
Agam   >GCCACCCAGAAGACGGTCGATGGCCCCTCGGGCAAGCTGTGGCGTGATGGCCGTGGTGCCGCC>  3
Bmor   >GCTACACAGAAAACTGTTGATGGACCTTCTGGAAAATTATGGCGTGATGGCCGTGGTGCT--->  2
Agla   >GCAACTCAGAAGACTGTGGACGGACCTTCTGGAAAATTGTGGCGTGATGGTCGTGGCGCTGGA>  5
        ..|..|..|..|..|..|..|..|..|..|..|..|..|..|..|..|..|..|..|..|..|

         Q  N  I  I  P  A  x  T  G  A  A  K  A  V  G  K  V  I  P  A  L
Dmel1  >CAGAACATCATCCCGGCCGCCACCGGAGCCGCCAAGGCTGTGGGCAAGGTCATCCCCGCCCTG>  1
Dmel2  >CAGAACATCATTCCAGCTTCCACTGGAGCTGCCAAGGCCGTGGGCAAGGTTATCCCCGCCCTC>  2
Mdom   >CAAAACATCATTCCTGCTTCCACCGGTGCCGCCAAGGCTGTCGGCAAAGTTATTCCTGCCTTG>  0
Scal   >CAAAATATCATTCCTGCCTCCACCGGTGCCGCCAAGGCTGTCGGCAAAGTAATTCCTGCCTTG>  2
Agam   >CAGAACATCATTCCGGCGGCGACCGGTGCCGCCAAGGCGGTGGGCAAGGTCATCCCGGCCCTG>  4
Bmor   >CAA/
        CAAAACATCATTCCTGCCTCTACTGGTGCTGCCAAAGCTGTGGGTAAGGTTATCCCTGCTCTT>  3
Agla   >CAAAACATCATTCCTGCCTCCACTGGAGCCGCTAAAGCAGTCGGGAAGGTCATTCCTGCCCTT>  3
        ..|..|..|..|..|..|..|..|..|..|..|..|..|..|..|..|..|..|..|..|..|

         N  G   K  L  T  G  M  A  F  R  V  P  x  x   N  V  S  V  V  D
Dmel1  >AACG GCAAGCTGACCGGCATGGCTTTCCGCGTGCCCACG-CCCAATGTCTCCGTTGTGGAT>  2
Dmel2  >AACG GTAAGCTCACCGGAATGGCATTCCGTGTGCCCACT-CCCAACGTTTCCGTGGTCGAT>  2
Mdom   >AACG GTAAACTCACTGGTATGGCTTTCCGTGTGCCCACT-CCCAATGTATCTGTTGTCGAT>  0
Scal   >AATG GCAAACTCACTGGCATGGCTTTCCGTGTGCCCACC-CCCAATGTGTCGGTTGTCGAT>  1
Agam   >AACG GCAAGCTGACCGGTATGGCGTTCCGCGTCCCGACC-CCGAACGTCTCGGTCGTCGAT>  4
Bmor   >AATG GCAAGCTGACTGGAATGGCATTCCGCGTCCCTGTT-GCTAATGTATCTGTTGTTGAT>  5
Agla   >AATG^GCAAACTTACTGGCATGGCTTTCCGTGTACC-ACTAGCCAATGTATCTGTTGTCGAT>  2
        ..|. .|..|..|..|..|..|..|..|..|..|..|..|..| ..|..|..|..|..|..|..|

         L  T  V  R  L  x  K  x  A  x  Y  x  x  I  K  x  K  V
Dmel1  >CTTACCGTCCGCTTGGGCAAGGGAGCCACCTATGACGAAATCAAGGCTAAGGTC>  1
Dmel2  >TTGACCGTGCGCTTGGGCAAGGGTGCGTCCTATGATGAAATTAAGGCCAAGGTT>  2
Mdom   >TTGACTGTCCGTCTTGGCAAGCCAGCCAACTACGAAGCCATCAAAGCCAAGGTT>  0
Scal   >TTGACTGTCCGCCTTGGCAAGCCAGCCAACTATGAAGCCATCAAAGCCAAGGTT>  1
Agam   >CTGACCGTGCGCCTGTCCAAGCCGGCCACCTACGACCAGATCAA-GC--AG/
        ................................................AAGGTG>  5
Bmor   >CTAACTGTTCGTCTTGGAAAACCTGCAAGCTATGAAGCCATCAA-GC-AA/
        ................................................AAGGTC>  5
Agla   >CTGACAGTACGTCTGGGCAAAGGGGCCTCGTACGATGAAATCAAAGCCAAAGTT>  4
        ..|..|..|..|..|..|..|..|..|..|..|..|..|..|..|..|..|..|

         x  E  A  x  x  G  x  x  K  G  I  L  x  Y  T  x  E  E  V  V  S
Dmel1  >GAGGAGGCCTCCAAGGGACCCCTGAAGGGAATCCTGGGCTACACCGATGAGGAGGTGGTCTCC>  2
Dmel2  >CAGGAGGCCGCCAACGGACCCCTGAAGGGTATCCTGGGATACACCGATGAGGAGGTCGTTTCT>  2
Mdom   >TTGGAGGCTTCCAATGGACCCATGAAGGGTATCTTGGGCTACACCGATGAAGAAGTCGTCTCC>  1
Scal   >TTGGAAGCTTCCAACGGTCCCATGAAGGGCATCTTGGGCTACACCGATGAGGAAGTAGTCTCC>  2
Agam   >AAGGAGGCCGCCAACGGGCCGATGAAGGGCATCCTGGACTACACCGAGGAGGAGGTCGTGTCG>  5
Bmor   >AAGGAGGCAGCTGAAGGTCCTTTGAAGGGCATTCTCGGGTATACTGAAGATCAAGTTGTGTCC> 12
Agla   >AAGGAGGCCGCCGAGGGTGACCTCAAGGGAATCTTGGGTTACACAGATGAAGAAGTTGTCTCA>  5
        ..|..|..|..|..|..|..|..|..|..|..|..|..|..|..|..|..|..|..|..|..|
```

**Fig. S2 cont.** (part 4 of 4)

```
         x  D  F  x  x  D  x  H  S  S  x  F  D  A  x  A  G  I  x  L
Dmel1  >ACCGACTTCTTCAGCGACACCCATTCGTCTGTGTTCGACGCCAAGGCTGGCATTTCGCTG>  1
Dmel2  >ACCGATTTCCTCAGCGACACCCACTCGTCGGTGTTCGATGCCAAGGCTGGCATTTCGCTA>  2
Mdom   >ACTGACTTCGTCAGCGACACCCACTCCTCGGTCTTCGATGCCAAGGCCGGTATCTCCCTC>  3
Scal   >ACCGACTTCGTCAGCGATACCCACTCCTCTGTGTTCGACGCCAAGGCTGGCATCTCCCTC>  0
Agam   >ACCGACTTCGTCGGCGACTGCCACTCGTCCATCTTTGACGCGAAGGCTGGCATCCAGCTG>  4
Bmor   >TCAGACTTCATTGGTGATTCACACTCTTCAATCTTTGATGCTGCCGCTGGAATTTCTTTG>  6
Agla   >TCCGATTTCATTGGTGACACACATTCTTCCATCTTTGACGCAAAGGCTGGAATTTCTTTG>  1
         ..|..|..|..|..|..|..|..|..|..|..|..|..|..|..|..|..|..|..|..|

         x  x  x  F  V  K  L  I  S  W  Y  D  N  E  x  G  Y
Dmel1  >AACGATAAGTTCGTCAAGCTAATCTCGTGGTACGACAACGAGTTCGGTTAC>  2
Dmel2  >AACGACAAGTTCGTGAAGCTGATCTCTTGGTACGACAACGAGTTTGGCTAC>  0
Mdom   >AACGACAACTTCGTCAAGTTGATCTCCTGGTACGACAACGAATTCGGTTAC>  0
Scal   >AACGAT/
         .....CAA-TTTGTCAAGTTGATCTCCTGGTACGACAACGAATTCGGTTAC>  0
Agam   >AGCGACACGTTCGTCAAGCTGATCTCCTGGTACGACAACGAGTACGGCTAC>  2
Bmor   >AATGACAACTTTGTGAAGCTGATCAGTTGGTACGACAATGAATATGGTTAT>  3
Agla   >AATCAAAACTTCGTCAAACTCATCTCTTGGTATGACAATGAATACGGTTAC>  5
         ..|..|..|..|..|..|..|..|..|..|..|..|..|..|..|..|..|

         S  x  R  V  x  D  L  I  K  Y  x  x  x  K  D  x  *
Dmel1  >TCCAACCGCGTCATCGACCTGATCAAGTATATGCAGAGCAAGGAC---TAA>  2   0.478 ( 9/ 19)
Dmel2  >TCCAACCGCGTCATCGACCTGATCAAGTACATGCAGAGCAAGGAT---TAA>  0   0.062 ( 1/ 16)
Mdom   >TCCAACCGTGTCATTGACTTGATCAAATACATGCAATCCAAGGAT---TAA>  0   0.000 ( 0/  9)
Scal   >TCCAACCGTGTCATCGACTTGATCAAATACATGCAATCCAAGGAT---TAA>  0   0.235 ( 4/ 17)
Agam   >TCGAACCGCGTCGTCGATCTGATCAAGTACATGCAGACCAAGGAT---TAA>  2   0.227 (15/ 66)
Bmor   >TCCAGCAGAGTCATTGATCTCATCAAGTACATTCAATCTAAAGAT---TAA>  3   0.322 (28/ 87)
Agla   >TCCACCAGGGTCATCGACTTGATTAAATACATTTCCACTAAAGATGCCTAA>  6   0.188 (16/ 85)
         ..|..|..|..|..|..|..|..|..|..|..|..|..|..|..|..|..|      0.266 (57/214)
```

Fig. S2 continued. Expected unique differences given TR content (see text)

Dmel1 289/999 = 0.289   0.289(19) = **5.5 substitutions predicted, 9 observed**
CGCGCCGCCGCGCGTGGTGGTCGACTCGACTCTGGTGGTGAGAAGAAGCGAGCGCGGTGGTGGGTGTTCACCACCAGGCGGCGCAAGAA
GTCATCATCTCCCGCCGGCGCGTGTGCGGCGGGTGGTCTCCACCACCATCATCAGAGATCGTCGTGATGACCACCCCACCACAGAAGAC
TCTTGTGGCGCGCGCCGCCCATCATCGCCGCCGCCGCCTGTGCGCGTCTCTGTGGAGGAGGACACGAGGAGGTGGTCTCTTCTTCACAC
TGTGTTCTCCGCGTATATAGAG

Dmel2 244/999 = 0.244   0.244(16) = **3.9 substitutions predicted, 1 observed**
TCTCCGCGCCGCCGCGCTGTGTGTGTTCGATTCGAGCCGCCGTGGTGGTAGAAGACGCGGCGCGTGTTCACCACCACACTCACTAAGAA
GATCATCTCCCGCCGGCGGCGGGTGGTCTCCACCACCACTCCTCTGAGATCGTCGTGATGACCACCCCACCACCAGAAGATGTGCATCA
TGTGTGGCGCATGATGAAGGAGGCCGCCACACGAGGAGGACACTCGTCGGTGTTCTCTCGCGAGAG

Mdom  178/999 = 0.178   0.178(9) = **1.6 substitution predicted, 0 observed**
GCCGCCGCGCGCGCATATGTTGTTGGTGTCACCACCAGGTGGTGAAGAAGTCATCATCTCCTGCTGGTGTGGTGTTGTGCGCGCACCAC
CATCTCCCACCACTGTGCATCATGCCGCCGTGTGTGTTGTAGCCAGCCAACACGAAGAAGTCGTCTCACACCTCCTCTCTCTCTCGTGT

Scal  218/999 = 0.218   0.218(17) = **3.7 substitutions predicted, 4 observed**
GCGCTGTGTGTGATTCGATTCGATGCTGCGTTGTTCGCGGCGCATATGTTGTGCACCACCAGGTGGTGAAGAAGGTCATCATCTCTGTG
TTCTCCCTCCTGCACCACCATTGTTGTCTCTGTGACTACTCACACACTCTTGTGGTGGTGCTGCATATGCCGCCGTGTGTGTGTAGCCA
GCCAACACAGTAGTCTCCTCCTCTGTGTTCTCTCTCGTGT

Agam  289/999 = 0.289  0.289(66) = **19.0 substitutions predicted, 15 observed**
GTCGTCTGCTGCGCGCCGCCGGTGGTGTGTGTGTAGAAGAGTGTGCGCGGCGCGCGTCGTGTGTGTCACCACGAGAGCGCAAGAAGTCA
TCATCTCCGCCGATGCGCCGATGTCGTCGTCGGTGTGTGTCGCGCACCACCATCTCTCATCATGATGACGACGCGCGACCACCAGAAGA
TGTGGTGGTGCCGCCCATCATCGGCGGCGGCCGCCCGCGTCTCGTCGTCGGCGCAGGAGGCCGCCACACGAGGAGGAGGGTGTTCGTCG
CGCGACACTCTCCGCGTCGTCG

Bmor   230/999 = 0.230  0.230(87) = **20.0 substitutions predicted, 28 observed**
TATATATATGTGTGGATGGATTTGTTGTTATATGTTGTAGAGGTGTGGAGGAGGAAAAAATTATTATATTGTGGTGTCTCTTCTCCTT
CTTCACCACATTGTTGTGATGACTACTCACAACACAGTGGTGCAACAACATCATCTCTTGCTGCTGTGCTCTCGCGTGTTGTTGAGGAG
GTCTCTATATGTGTCACACTCTTGCTGCTGTGATATCAGCAGAGTCTCATCA

All 1448/5994 = 0.242   0.242(217) = **52.4 substitutions predicted, 57 observed**
CGCGCCGCCGCGCGTGGTGGTCGACTCGACTCTGGTGGTGAGAAGAAGCGAGCGCGGTGGTGGGTGTTCACCACCAGGCGGCGCAAGAA
GTCATCATCTCCCGCCGGCGCGTGTGCGGCGGGTGGTCTCCACCACCATCATCAGAGATCGTCGTGATGACCACCCCACCACAGAAGAC
TCTTGTGGCGCGCGCCGCCCATCATCGCCGCCGCCGCCTGTGCGCGTCTCTGTGGAGGAGGACACGAGGAGGTGGTCTCTTCTTCACAC
TGTGTTCTCCGCGTATATAGAGTCTCCGCGCCGCCGCGCTGTGTGTGTTCGATTCGAGCCGCCGTGGTGGTAGAAGACGCGGCGCGTGT
TCACCACCACACTCACTAAGAAGATCATCTCCCGCCGGCGGCGGGTGGTCTCCACCACCACTCCTCTGAGATCGTCGTGATGACCACCC
CACCACCAGAAGATGTGCATCATGTGTGGCGCATGATGAAGGAGGCCGCCACACGAGGAGGACACTCGTCGGTGTTCTCTCGCGAGAGG
CCGCCGCGCGCGCATATGTTGTTGGTGTCACCACCAGGTGGTGAAGAAGTCATCATCTCCTGCTGGTGTGGTGTTGTGCGCGCACCACC
ATCTCCCACCACTGTGCATCATGCCGCCGTGTGTGTTGTAGCCAGCCAACACGAAGAAGTCGTCTCACACCTCCTCTCTCTCTCGTGTG
CGCTGTGTGTGATTCGATTCGATGCTGCGTTGTTCGCGGCGCATATGTTGTGCACCACCAGGTGGTGAAGAAGGTCATCATCTCTGTGT
TCTCCCTCCTGCACCACCATTGTTGTCTCTGTGACTACTCACACACTCTTGTGGTGGTGCTGCATATGCCGCCGTGTGTGTGTAGCCAG
CCAACACAGTAGTCTCCTCCTCTGTGTTCTCTCTCGTGTGTCGTCTGCTGCGCGCCGCCGGTGGTGTGTGTGTAGAAGAGTGTGCGCGG
CGCGCGTCGTGTGTGTCACCACGAGAGCGCAAGAAGTCATCATCTCCGCCGATGCGCCGATGTCGTCGTCGGTGTGTGTCGCGCACCAC
CATCTCTCATCATGATGACGACGCGCGACCACCAGAAGATGTGGTGGTGCCGCCCATCATCGGCGGCGGCCGCCCGCGTCTCGTCGTCG
GCGCAGGAGGCCGCCACACGAGGAGGAGGGTGTTCGTCGCGCGACACTCTCCGCGTCGTCGTATATATATGTGTGGATGGATTTGTTGT
TATATGTTGTAGAGGTGTGGAGGAGGAAAAAATTATTATATTGTGGTGTCTCTTCTCCTTCTTCACCACATTGTTGTGATGACTACTC
ACAACACAGTGGTGCAACAACATCATCTCTTGCTGCTGTGCTCTCGCGTGTTGTTGAGGAGGTCTCTATATGTGTCACACTCTTGCTGC
TGTGATATCAGCAGAGTCTCATCA

**Figure S3. Gapdh MSA with locations of triplet nucleotide repeats.** There are 33 unique positions where an amino acid is repeated in the Gapdh protein MSA. Those shared by more than one lineage are highlighted in yellow, while those found in a single lineage (10/33 sites) are highlighted in cyan. Underlined regions are regions corresponding to trinucleotide repeats. Change in the underlining style (bold or double underlining) is used to contrast with adjacent repeats and to indicate places where the tandem repeat is out of frame with the codon frame (double underlining).

```
                                     1       2                                      3
D_mel_Gapdh1        MSKIGINGFGRIGRLVLRAAIDKGASVVAVNDPFIDVNYMVYLFKFDSTHGRFKGTVAAE
D_mel_Gapdh2        MSKIGINGFGRIGRLVLRAAIDKGANVVAVNDPFIDVNYMVYLFKFDSTHGRFKGTVAAE
M_dom_Gapdh         MSKIGINGFGRIGRLVLRAAIDKGAQVVAVNDPFIDVNYMVYLFKFDSTHGRFKGTVAAE
S_cal_Gapdh         MSKIGINGFGRIGRLVLRAAIDKGAQVVAVNDPFIDVEYMVYLFKFDSTHGRFKGTVAAE
A_gam_Gapdh         MSKIGINGFGRIGRLVLRAAITKGASVVAINDPFIGVDYMVYLFKYDSTHGRFKGEVSAQ
B_mor_Gapdh         MSKIGINGFGRIGRLVLRASIEKGAQVVAINDPFIGLDYMVYLFKYDSTHGRFKGSVEVQ
A_gla_Gapdh         MVKIGINGFGRIGRLVLRASIERGAEVVAVNDPFLDVDYMVYLFKFDSTHGRYKGCVSSD

                     4   5   6                       7       8           9 10 11
D_mel_Gapdh1        GGFLVVNGQKITVFSERDPANINWASAGAEYVVESTGVFTTIDKASTHLKGGAKKVIISA
D_mel_Gapdh2        GGFLVVNGQKITVFSERDPANINWASAGAEYIVESTGVFTTIDKASTHLKGGAKKVIISA
M_dom_Gapdh         GGKLVVNGQAITVFSERDPANINWASAGAEYVVESTGVFTTIDKASAHFKGGAKKVIISA
S_cal_Gapdh         GGKLVVNGQAITVFSERDPANINWASAGAEYVVESTGVFTTIDKASAHFKGGAKKVIISA
A_gam_Gapdh         DGCLVVNGQKIAVFQERDPKAIPWGKAGAEYVVESTGVFTTTEKASAHLEGGAKKVIISA
B_mor_Gapdh         DGFLVVNGNKIAVFSERDPKAIPWGKAGAEYVVESTGVFTTTDKASAHLEGGAKKVIISA
A_gla_Gapdh         GKNLVVDGKKIAVFQERDPKAIPWGQSGADYVVESTGVFTTIEKASAHLAGGAKKVIISA

                                12             13      14                   15    16
D_mel_Gapdh1        PSADAPMFVCGVNLDAYSPDMKVVSNASCTTNCLAPLAKVINDNFEIVEGLMTTVHATTA
D_mel_Gapdh2        PSADAPMFVCGVNLDAYKPDMKVVSNASCTTNCLAPLAKVINDNFEIVEGLMTTVHATTA
M_dom_Gapdh         PSADAPMFVCGVNLDAYKPDMKVVSNASCTTNCLAPLAKVIHDNFEIVEGLMTTVHATTA
S_cal_Gapdh         PSADAPMFVCGVNLDAYKPDMKVVSNASCTTNCLAPLAKVIHDNWEIVEGLMTTVHATTA
A_gam_Gapdh         PSADAPMFVVGVNLEAYEPSMKVVSNASCTTNCLAPLAKVINDNFGILEGLMTTVHATTA
B_mor_Gapdh         PSADAPMFVVGVNLEAYDPSFKVISNASCTTNCLAPLAKVIHDNFEIVEGLMTTVHATTA
A_gla_Gapdh         PSADAPMYVCGVNLDAYNPADKVISNASCTTNCLAPLAKVIHDNFEIVEGLMTTVHATTA

                                   17/18 19 20   21                                22
D_mel_Gapdh1        TQKTVDGPSGKLWRDGRGAAQNIIPAATGAAKAVGKVIPALNGKLTGMAFRVPTPNVSVV
D_mel_Gapdh2        TQKTVDGPSGKLWRDGRGAAQNIIPASTGAAKAVGKVIPALNGKLTGMAFRVPTPNVSVV
M_dom_Gapdh         TQKTVDGPSGKLWRDGRGAAQNIIPASTGAAKAVGKVIPALNGKLTGMAFRVPTPNVSVV
S_cal_Gapdh         TQKTVDGPSGKLWRDGRGAAQNIIPASTGAAKAVGKVIPALNGKLTGMAFRVPTPNVSVV
A_gam_Gapdh         TQKTVDGPSGKLWRDGRGAAQNIIPAATGAAKAVGKVIPALNGKLTGMAFRVPTPNVSVV
B_mor_Gapdh         TQKTVDGPSGKLWRDGRGAQQNIIPASTGAAKAVGKVIPALNGKLTGMAFRVPVANVSVV
A_gla_Gapdh         TQKTVDGPSGKLWRDGRGAGQNIIPASTGAAKAVGKVIPALNGKLTGMAFRVPLANVSVV

                                         2324            25262728 29    30   31
D_mel_Gapdh1        DLTVRLGKGATYDEIKAKVEEASKGPLKGILGYTDEEVVSTDFFSDTHSSVFDAKAGISL
D_mel_Gapdh2        DLTVRLGKGASYDEIKAKVQEAANGPLKGILGYTDEEVVSTDFLSDTHSSVFDAKAGISL
M_dom_Gapdh         DLTVRLGKPANYEAIKAKVLEASNGPMKGILGYTDEEVVSTDFVSDTHSSVFDAKAGISL
S_cal_Gapdh         DLTVRLGKPANYEAIKAKVLEASNGPMKGILGYTDEEVVSTDFVSDTHSSVFDAKAGISL
A_gam_Gapdh         DLTVRLSKPATYDQIKQKVKEAANGPMKGILDYTEEEVVSTDFVGDCHSSIFDAKAGIQL
B_mor_Gapdh         DLTVRLGKPASYEAIKQKVKEAAEGPLKGILGYTEDQVVSSDFIGDSHSSIFDAAAGISL
A_gla_Gapdh         DLTVRLGKGASYDEIKAKVKEAAEGDLKGILGYTDEEVVSSDFIGDTHSSIFDAKAGISL

                                   32 33
D_mel_Gapdh1        NDKFVKLISWYDNEFGYSNRVIDLIKYMQSKD-
D_mel_Gapdh2        NDKFVKLISWYDNEFGYSNRVIDLIKYMQSKD-
M_dom_Gapdh         NDNFVKLISWYDNEFGYSNRVIDLIKYMQSKD-
S_cal_Gapdh         NDQFVKLISWYDNEFGYSNRVIDLIKYMQSKD-
A_gam_Gapdh         SDTFVKLISWYDNEYGYSNRVVDLIKYMQTKD-
B_mor_Gapdh         NDNFVKLISWYDNEYGYSSRVIDLIKYIQSKD-
A_gla_Gapdh         NQNFVKLISWYDNEYGYSTRVIDLIKYISTKDA
```

**Figure S4. Point differences are enriched in TRs.** Point differences are shown between this pairwise comparison of *D. melanogaster gapdh1* and *A. gambiae gapdh*. Differences within perfect TRs are highlighted in yellow, while differences within imperfect repeats with transition mutations are highlighted in gray if not already highlighted in yellow. Point differences within lineage-specific extensions of shared TRs are counted as lineage-specific differences.

```
Dmel  ATGTCGAAGATCGGAATTAACGGATTTGGCCGCATCGGCCGCTTGGTGCTCCGCGCCGCC
Agam  ATGTCGAAGATCGGAATTAACGGATTTGGCCGCATCGGTCGTCTGGTGCTGCGCGCCGCC
      ......................................Y..YY.......S.........

Dmel  ATCGATAAGGGCGCCTCCGTGGTGGCCGTCAACGATCCCTTCATCGATGTCAACTACATG
Agam  ATCACCAAGGGTGCGTCGGTGGTCGCCATCAACGATCCGTTCATCGGCGTCGACTACATG
      ...RMY.....Y..S..S.....S...R..........S.......RY...R........

Dmel  GTTTACCTGTTTAAATTCGACTCGACTCACGGTCGTTTCAAGGGCACCGTTGCGGCTGAG  alt. imperfect
Dmel  GTTTACCTGTTTAAATTCGACTCGACTCACGGTCGTTTCAAGGGCACCGTTGCGGCTGAG
Agam  GTGTACCTGTTCAAGTACGACTCGACGCACGGTCGCTTCAAGGGCGAGGTGTCCGCCCAG
      ..K........Y..R.W.........K........Y.........RMS..KK.S..YS..

Dmel  GGCGGATTCCTGGTGGTGAACGGCCAGAAGATCACCGTGTTCAGCGAGCGCGACCCGGCCAAC
Agam  GACGGCTGCCTGGTCGTGAACGGCCAGAAGATTGCCGTGTTCCAGGAGCGCGACCCGAAGGCG
      .R...M.K......S..................YR........MRS............RMSRMS

Dmel  ATCAACTGGGCCAGTGCTGGAGCCGAGTATGTGGTGGAGTCCACCGGAGTGTTCACCACCATT
Agam  ATCCCGTGGGGTAAGGCCGGCGCGGAGTACGTCGTTGAGTCGACCGGTGTGTTCACCACGACC
      ...MMS....SY.RK..Y..M..S.....Y..S..K.....S.....W...........S.YY

Dmel  GACAAGGCGTCCACCCACTTGAAGGGCGGCGCCAAGAAGGTCATCATCTCGGCCCCA
Agam  GAGAAGGCATCCGCCCATCTGGAGGGTGGCGCCAAGAAGGTCATCATCTCGGCCCCG
      ..S.....R...R....YY..R....Y.............................R

Dmel  TCCGCCGATGCGCCCATGTTCGTGTGCGGCGTTAACCTGGACGCCTACAGCCCCGACATG
Agam  TCCGCCGATGCGCCGATGTTCGTCGTCGGTGTCAACCTGGAGGCGTACGAGCCATCGATG
      ..............S.........SKK...Y..Y........S..S...RRS..MKMS...

Dmel  AAGGTGGTCTCCAACGCCTCGTGCACCACCAACTGCCTGGCTCCCCTGGCCAAG
Agam  AAGGTCGTGTCGAACGCGTCCTGCACCACCAACTGTCTCGCCCCGCTGGCCAAG
      .....S..S..S.....S..S..............Y..S..Y..S.........

Dmel  GTCATCAATGACAACTTCGAGATCGTCGAGGGTCTGATGACCACCGTGCACGCCACCACT
Agam  GTCATCAACGACAACTTCGGCATCCTGGAGGGTCTGATGACGACGGTGCACGCGACCACC
      ........Y..........RS...S.S..............S..S........S.....Y

Dmel  GCCACCCAGAAGACCGTCGACGGTCCCTCTGGCAAACTGTGGCGCGATGGACGTGGCGCCGCC
Agam  GCCACCCAGAAGACGGTCGATGGCCCCTCGGGCAAGCTGTGGCGTGATGGCCGTGGTGCCGCC
      ..............S.....Y..Y.....K.....R........Y.....M.....Y......

Dmel  CAGAACATCATCCCGGCCGCCACCGGAGCCGCCAAGGCTGTGGGCAAGGTCATCCCCGCCCTG
Agam  CAGAACATCATTCCGGCGGCGACCGGTGCCGCCAAGGCGGTGGGCAAGGTCATCCCGGCCCTG
      ...........Y.....S..S.....W...........K.................S......

Dmel  AACGGCAAGCTGACCGGCATGGCTTTCCGCGTGCCCACGCCCAATGTCTCCGTTGTGGAT
Agam  AACGGCAAGCTGACCGGTATGGCGTTCCGCGTCCCGACCCCGAACGTCTCGGTCGTCGAT
      .................Y.....K........S..S..S..S.Y......S..Y..S...
```

**Fig. S4 cont.** (part 2 of 2)

```
Dmel   CTTACCGTCCGCTTGGGCAAGGGAGCCACCTATGACGAAATCAAGGCTAAGGTC
Agam   CTGACCGTGCGCCTGTCCAAGCCGGCCACCTACGACCAGATCAAGCAGAAGGTG
       ..K.....S...Y..KS....SSR........Y...S.R......SMK.....S

Dmel   GAGGAGGCCTCCAAGGGACCCCTGAAGGGAATCCTGGGCTACACCGATGAGGAGGTGGTCTCC
Agam   AAGGAGGCCGCCAACGGGCCGATGAAGGGCATCCTGGACTACACCGAGGAGGAGGTCGTGTCG
       R.........K.....S..R..SM.......M.......R.........K.........S..S..S

Dmel   ACCGACTTCTTCAGCGACACCCATTCGTCTGTGTTCGACGCCAAGGCTGGCATTTCGCTG
Agam   ACCGACTTCGTCGGCGACTGCCACTCGTCCATCTTTGACGCGAAGGCTGGCATCCAGCTG
       ..........K..R.....WS...Y.....YR.S..Y.....S...........YYM....

Dmel   AACGATAAGTTCGTCAAGCTAATCTCGTGGTACGACAACGAGTTCGGTTAC
Agam   AGCGACACGTTCGTCAAGCTGATCTCCTGGTACGACAACGAGTACGGCTAC
       .R...Y.M...........R.....S................W...Y...

Dmel   TCCAACCGCGTCATCGACCTGATCAAGTATATGCAGAGCAAGGACTAA
Agam   TCGAACCGCGTCGTCGATCTGATCAAGTACATGCAGACCAAGGATTAA
       ..S.........R.....Y............Y.........S......Y...
```

| Difference | Type | Outside TRs | Inside TRs |
|---|---|---|---|
| Y = CT | transition | 25.3% (23/91) | 23.3% (21/90) |
| R = AG | transition | 23.1% (21/91) | 10.0% ( 9/90) |
| S = CG | transversion, CG-rich | 26.4% (24/91) | 46.7% (42/90) |
| K = GT | transversion | 7.7% ( 7/91) | 13.3% (12/90) |
| M = AC | transversion | 15.4% (14/91) | 3.3% ( 3/90) |
| W = AT | transversion, CG-poor | 2.2% ( 2/91) | 3.3% ( 3/90) |
| **transitions** | all transitions | **48.4% (44/91)** | **33.3% (30/90)** |
| **transversions** | all transversions | **51.6% (47/91)** | **66.7% (60/90)** |

**Figure S5. *Drosophila melanogaster Gapdh1* and *Gapdh2* arose from a melanogaster species group duplication.** Shown is a neighbor-joining tree depicting the duplication of *gapdh* into *Gapdh1* and *Gapdh2* in the stem-lineage of the melanogaster species group. Boot strap support values ≥ 50% from 500 boot strap replicates are shown. The tree is rooted on the obscura species group, which has a single gene. Other non-duplicated lineages in the tree correspond to the lineages from the montium and ananassae species groups.

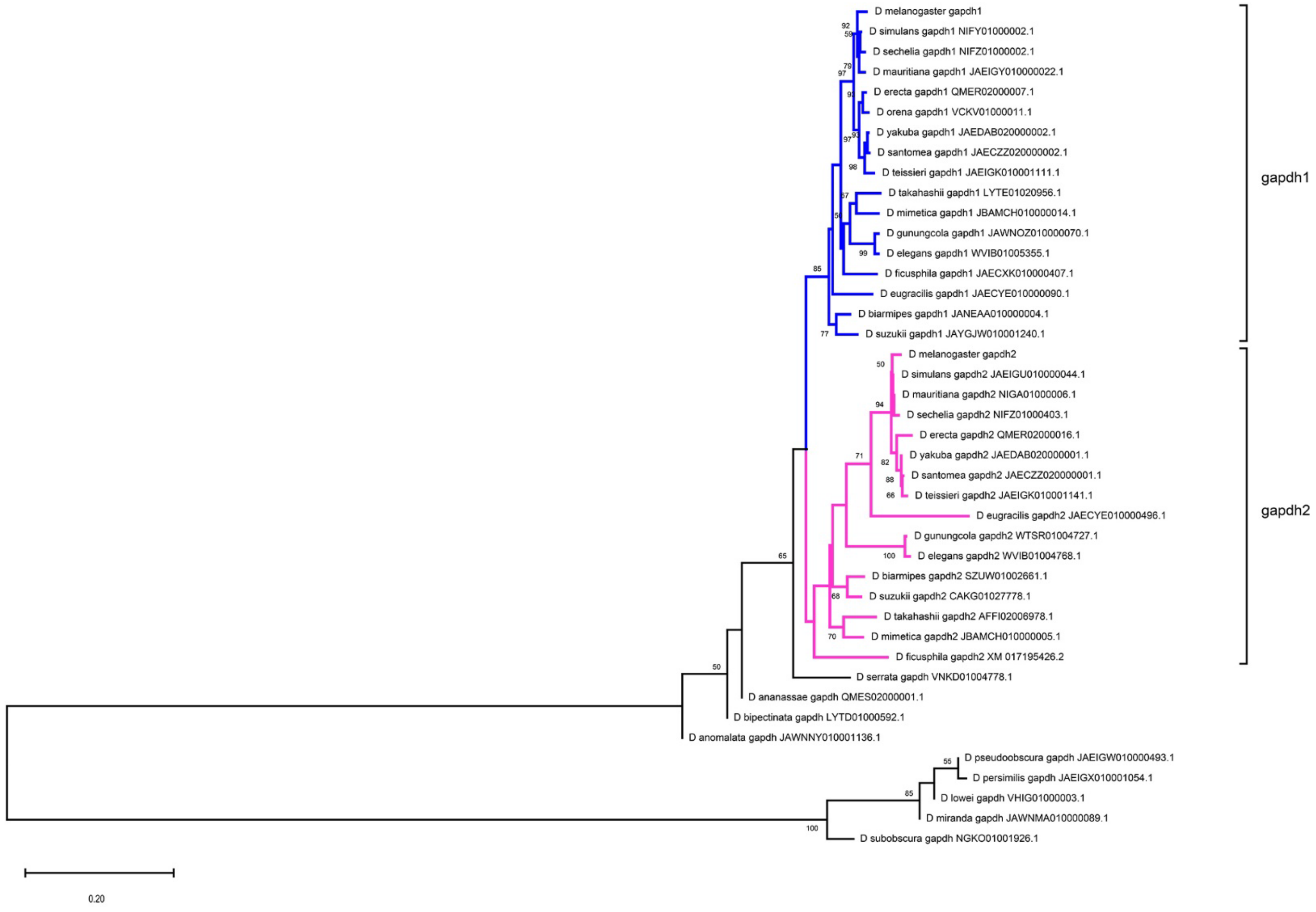

**Figure S6. *Drosophila melanogaster Gapdh1* polymorphisms.** Shown is an alignment of *Gapdh1* and 10 different SNPs identified in DGRP data. Half of these (5/10) are C↔T (Y) transitions that are not associated with de novo TR origination. The other half (5/10) are all associated with de novo repeat creation. Considered as RS events none of these would count as substitutions. Considered as substitutions, they include two G↔T (K) transversions and three transitions (two Y and one R). Residues highlighted in cyan are derived changes. Residues highlighted in gray are unpolarized differences detected in the outgroup clade. Number of DGRP lines with unambigious genotypes are indicated.

```
Gapdh1 SNP #1:            152 C
                          228 T
Dmela min  ATGTCGAAGATCGGAATCAACGGATTTGGCCGCATCGGCCGCTTGGTGCTCCGCGCCGCC
Dmela MAJ  ATGTCGAAGATCGGAATTAACGGATTTGGCCGCATCGGCCGCTTGGTGCTCCGCGCCGCC
Dsimu      ATGTCGAAGATCGGAATTAACGGATTTGGCCGCATCGGCCGCCTGGTGCTCCGCGCCGCC
Dsech      ATGTCGAAGATCGGAATTAACGGATTTGGCCGCATCGGCCGCCTGGTGCTCCGCGCCGCC
Dmaur      ATGTCGAAGATCGGAATTAACGGATTTGGCCGCATCGGCCGCCTGGTGCTCCGCGCCGCC
Derec      ATGTCGAAGATCGGAATTAACGGATTTGGCCGCATCGGCCGTCTGGTGCTCCGCGCCGCC
Doren      ATGTCGAAGATCGGAATTAACGGATTTGGCCGCATCGGCCGTCTGGTGCTCCGCGCCGCC
Dyaku      ATGTCGAAGATTGGAATTAACGGATTTGGCCGCATCGGCCGTCTGGTGCTCCGCGCCGCC
Dsant      ATGTCGAAGATTGGAATTAACGGATTTGGCCGCATCGGCCGTCTGGTGCTCCGCGCCGCC
```

```
Gapdh1 SNP #2:                38 T
                             354 G
Dmela min  ATCGATAAGGGCGCCTCCGTTGTGGCCGTCAACGATCCCTTCATCGATGTCAACTACATG
Dmela MAJ  ATCGATAAGGGCGCCTCCGTGGTGGCCGTCAACGATCCCTTCATCGATGTCAACTACATG
Dsimu      ATCGATAAGGGCGCCTCCGTGGTGGCCGTCAACGATCCCTTCATCGATGTCAACTACATG
Dsech      ATCGATAAGGGCGCCTCCGTGGTGGCCGTCAACGATCCTTTCATCGATGTCAACTACATG
Dmaur      ATCGATAAGGGCGCCTCCGTGGTGGCCGTCAACGATCCCTTCATCGATGTCAACTACATG
Derec      ATCGATAAGGGCGCCTCCGTGGTGGCCGTCAACGATCCCTTCATCGATGTCAACTACATG
Doren      ATCGATAAGGGCGCCTCCGTGGTGGCCGTCAACGATCCCTTCATCGATGTCAACTACATG
Dyaku      ATCGATAAGGGCGCCTCTGTGGTGGCCGTCAACGATCCCTTCATCGATGTCAACTACATG
Dsant      ATCGATAAGGGCGCCTCTGTGGTGGCCGTCAACGATCCCTTCATCGATGTCAACTACATG


Dmela      GTTTACCTGTTTAAATTCGACTCGACTCACGGTCGTTTCAAGGGCACCGTTGCGGCTGAG
Dsimu      GTGTACCTGTTCAAATTCGACTCGACTCACGGTCGTTTCAAGGGCACCGTTGCGGCTGAG
Dsech      GTGTACCTGTTCAAATTCGACTCGACTCACGGTCGTTTCAAGGGCACCGTTGCGGCTGAG
Dmaur      GTGTACCTGTTCAAATTCGACTCGACTCACGGTCGTTTCAAGGGCACCGTTGCGGCTGAG
Derec      GTGTATCTGTTCAAATTCGACTCGACTCACGGTCGTTTCAAGGGCACCGTTGCGGCTGAG
Doren      GTGTATCTGTTCAAATTCGACTCGACTCACGGTCGTTTCAAGGGCACCGTTGCGGCTGAG
Dyaku      GTGTATCTCTTCAAATTCGACTCGACTCACGGTCGTTTCAAGGGCACCGTTGCGGCTGAG
Dsant      GTGTATCTCTTCAAATTCGACTCGACTCACGGTCGTTTCAAGGGCACCGTTGCGGCTGAG
```

```
Gapdh1 SNP #3:                         2 A
                                     404 G
Dmela min  GGCGGATTCCTGGTGGTGAACGGCCAGAAAATCACCGTGTTCAGCGAGCGCGACCCGGCC
Dmela MAJ  GGCGGATTCCTGGTGGTGAACGGCCAGAAGATCACCGTGTTCAGCGAGCGCGACCCGGCC
Dsimu      GGCGGATTCCTGGTGGTGAACGGCCAGAAGATCACCGTGTTCAGCGAGCGCGACCCGGCC
Dsech      GGCGGATTCCTGGTGGTGAACGGCCAGAAGATCACCGTGTTCAGCGAGCGCGACCCGGCC
Dmaur      GGCGGATTCCTGGTGGTGAACGGCCAGAAGATCACCGTGTTCAGCGAGCGCGACCCGGCC
Derec      GGCGGATTCCTGGTGGTGAACGGCCAGAAGATCACCGTGTTCAGCGAGCGCGACCCGGCC
Doren      GGCGGATTCCTGGTGGTGAACGGCCAGAAGATCACCGTGTTCAGCGAGCGCGACCCGGCC
Dyaku      GGCGGATTCCTGGTGGTGAACGGCCAGAAGATCACCGTGTTCAGCGAACGCGACCCCGCC
Dsant      GGCGGATTCCTGGTGGTGAACGGCCAGAAGATCACCGTGTTCAGCGAACGCGACCCCGCC
```

```
Gapdh1 SNP #4:                                                      4 T
                                                                  404 C
Dmela min  AACATCAACTGGGCCAGTGCTGGAGCCGAGTATGTGGTGGAGTCCACCGGAGTGTTCACT
Dmela MAJ  AACATCAACTGGGCCAGTGCTGGAGCCGAGTATGTGGTGGAGTCCACCGGAGTGTTCACC
Dsimu      AACATCAACTGGGCCAGTGCTGGAGCCGAGTATGTGGTGGAGTCCACCGGTGTGTTCACC
Dsech      AACATCAACTGGGCCAGTGCTGGAGCCGAGTATGTGGTGGAGTCCACCGGAGTGTTCACC
Dmaur      AACATCAACTGGGCCAGTGCTGGAGCCGAGTATGTGGTGGAGTCCACCGGAGTGTTCACC
Derec      AACATCAACTGGGCCAGCGCTGGAGCCGAGTATGTGGTGGAGTCCACCGGAGTGTTCACC
Doren      AACATCAACTGGGCCAGCGCTGGAGCCGAGTATGTGGTGGAGTCCACCGGTGTGTTCACC
Dyaku      AACATCAACTGGGCCAGCGCTGGAGCCGAGTATGTGGTGGAGTCCACCGGAGTGTTCACC
Dsant      AACATCAACTGGGCCAGCGCTGGAGCCGAGTATGTGGTGGAGTCCACCGGAGTGTTCACC

Gapdh1 (haplotypes not phased)
SNPs #5-7:  166 T                          170 C                    2 T
            210 C                          190 T                  398 G
Dmela min  ACCATTGACAAGGCGTCCACCCACTTGAAGGGCGGCGCCAAGAAGGTCATCATCTCTGCC
Dmela MAJ  ACCATCGACAAGGCGTCCACCCACTTGAAGGGCGGTGCCAAGAAGGTCATCATCTCGGCC
Dsimu      ACCATCGACAAGGCATCCACCCACTTGAAGGGCGGTGCCAAGAAGGTCATCATCTCGGCC
Dsech      ACCATCGACAAGGCATCCACCCACTTGAAGGGCGGTGCCAAGAAGGTCATCATCTCGGCC
Dmaur      ACCATCGACAAGGCGTCCACCCACTTGAAGGGCGGTGCCAAGAAGGTCATCATCTCGGCC
Derec      ACCATCGACAAGGCGTCCACCCACTTGAAGGGCGGTGCCAAGAAGGTCATCATCTCGGCC
Doren      ACCATCGACAAGGCTTCCACCCACTTGAAGGGCGGTGCCAAGAAGGTCATCATCTCGGCT
Dyaku      ACCATCGACAAGGCGTCCACCCACTTGAAGGGCGGTGCCAAGAAGGTCATCATCTCGGCC
Dsant      ACCATCGACAAGGCGTCCACCCACTTGAAGGGCGGTGCCAAGAAGGTCATCATCTCGGCC

Dmela      CCATCCGCCGATGCGCCCATGTTCGTGTGCGGCGTTAACCTGGACGCCTACAGCCCCGAC
Dsimu      CCATCCGCCGATGCGCCCATGTTCGTGTGCGGCGTTAACCTGGACGCCTACAGCCCCGAC
Dsech      CCATCCGCCGATGCGCCCATGTTCGTGTGCGGCGTTAACCTGGACGCCTACAGCCCCGAC
Dmaur      CCATCCGCCGATGCGCCCATGTTCGTGTGCGGCGTTAACCTGGACGCCTACAGCCCTGAC
Derec      CCATCCGCCGATGCGCCCATGTTCGTGTGCGGCGTGAACCTGGACGCCTACAGCCCCGAC
Doren      CCATCCGCCGATGCGCCCATGTTCGTGTGCGGCGTAAACCTGGACGCCTACAGCCCCGAC
Dyaku      CCATCCGCCGATGCGCCCATGTTCGTGTGCGGTGTGAACCTGGACGCCTACAGCCCCGAC
Dsant      CCATCCGCCGATGCGCCCATGTTCGTGTGCGGTGTAAACCTGGACGCCTACAGCCCCGAC

Dmela      ATGAAGGTGGTCTCCAACGCCTCGTGCACCACCAACTGCCTGGCTCCCCTGGCCAAGGTC
Dsimu      ATGAAGGTGGTCTCCAACGCCTCGTGCACCACCAACTGCCTGGCTCCCCTGGCCAAGGTC
Dsech      ATGAAGGTGGTCTCCAACGCCTCCTGCACCACCAACTGCCTGGCTCCCCTGGCCAAGGTT
Dmaur      ATGAAGGTGGTCTCCAACGCCTCCTGCACCACCAACTGCCTGGCTCCTCTGGCCAAGGTC
Derec      ATGAAGGTGGTCTCCAACGCCTCTTGCACCACCAACTGCCTGGCTCCCCTGGCTAAGGTC
Doren      ATGAAGGTGGTCTCCAACGCCTCTTGCACCACCAACTGCCTGGCTCCCCTGGCCAAGGTC
Dyaku      ATGAAGGTGGTCTCCAACGCCTCGTGCACCACCAACTGCCTGGCTCCCCTGGCCAAGGTC
Dsant      ATGAAGGTGGTCTCCAACGCCTCGTGCACCACCAACTGCCTGGCTCCCCTGGCCAAGGTC

Dmela      ATCAATGACAACTTCGAGATCGTCGAGGGTCTGATGACCACCGTGCACGCCACCACTGCC
Dsimu      ATCAATGACAACTTCGAGATCGTCGAGGGTCTGATGACCACCGTGCACGCCACCACTGCC
Dsech      ATTAATGACAACTTCGAGATCGTCGAGGGTCTGATGACCACCGTGCACGCCACCACTGCC
Dmaur      ATCAATGACAACTTCGAGATCGTCGAGGGTCTGATGACCACCGTGCACGCCACCACTGCC
Derec      ATCAATGACAACTTCGAGATCGTCGAGGGTCTGATGACCACCGTGCACGCCACCACTGCC
Doren      ATCAATGACAACTTCGAGATCGTCGAGGGTCTGATGACCACCGTGCACGCCACCACTGCC
Dyaku      ATCAATGACAACTTCGAGATCGTCGAGGGTCTGATGACCACCGTGCACGCCACCACTGCC
Dsant      ATCAATGACAACTTCGAGATCGTCGAGGGTCTGATGACCACCGTGCACGCCACCACTGCC
```

```
Dmela      ACCCAGAAGACCGTCGACGGTCCCTCTGGCAAACTGTGGCGCGATGGACGTGGCGCCGCC
Dsimu      ACCCAGAAGACCGTCGACGGTCCCTCCGGCAAACTGTGGCGCGATGGACGTGGCGCCGCC
Dsech      ACCCAAAAGACCGTCGACGGTCCCTCCGGCAAACTGTGGCGCGATGGACGTGGCGCCGCC
Dmaur      ACCCAGAAGACCGTCGACGGTCCCTCTGGCAAACTGTGGCGCGATGGACGTGGTGCCGCT
Derec      ACCCAGAAGACCGTCGACGGTCCCTCTGGCAAACTGTGGCGCGATGGACGTGGCGCCGCC
Doren      ACCCAGAAGACCGTCGACGGCCCCTCTGGCAAACTGTGGCGCGATGGACGTGGCGCCGCC
Dyaku      ACCCAGAAGACCGTCGACGGTCCCTCTGGCAAACTGTGGCGCGATGGACGTGGCGCCGCC
Dsant      ACCCAGAAGACCGTCGACGGTCCCTCTGGCAAACTGTGGCGCGATGGACGTGGCGCCGCC

Dmela      CAGAACATCATCCCGGCCGCCACCGGAGCCGCCAAGGCTGTGGGCAAGGTCATCCCCGCC
Dsimu      CAGAACATCATCCCGGCCGCCACCGGAGCCGCCAAGGCTGTGGGCAAGGTCATCCCGGCC
Dsech      CAGAACATCATCCCGGCTGCCACCGGAGCCGCCAAGGCTGTGGGCAAGGTCATCCCGGCC
Dmaur      CAGAACATCATCCCGGCCGCCACCGGAGCCGCCAAGGCTGTGGGCAAGGTCATCCCGGCC
Derec      CAAAACATCATCCCGGCCGCCACCGGAGCCGCCAAGGCTGTGGGCAAGGTCATCCCTGCC
Doren      CAAAACATCATCCCGGCCGCCACCGGAGCCGCCAAGGCTGTGGGCAAGGTCATCCCTGCC
Dyaku      CAGAACATCATCCCGGCCGCCACCGGAGCCGCCAAGGCTGTGGGCAAGGTCATCCCGGCC
Dsant      CAGAACATCATCCCGGCCGCCACCGGAGCCGCCAAGGCTGTGGGCAAGGTGATCCCGGCC

Dmela      CTGAACGGCAAGCTGACCGGCATGGCTTTCCGCGTGCCCACGCCCAATGTCTCCGTTGTG
Dsimu      CTGAACGGCAAGCTGACCGGCATGGCTTTCCGCGTGCCCACGCCCAATGTCTCCGTGGTG
Dsech      CTGAACGGCAAGCTGACCGGCATGGCTTTCCGCGTGCCCACGCCCAATGTCTCCGTGGTG
Dmaur      CTGAACGGCAAGCTGACCGGCATGGCTTTCCGCGTGCCCACGCCCAATGTCTCCGTGGTG
Derec      TTGAATGGAAAGCTGACCGGCATGGCTTTCCGCGTTCCCACTCCCAATGTCTCCGTGGTG
Doren      CTGAATGGAAAGCTGACCGGCATGGCTTTCCGCGTTCCCACTCCCAATGTCTCCGTGGTG
Dyaku      CTGAATGGAAAGCTGACCGGCATGGCTTTCCGCGTTCCCACTCCCAATGTCTCCGTCGTG
Dsant      CTGAATGGAAAGCTGACCGGCATGGCTTTCCGCGTTCCCACTCCCAATGTCTCCGTGGTG
```

***Gapdh1*** **SNP #8:**

```
                    162 T
                    214 C
Dmela min  GATCTTACCGTCCGCTTGGGCAAGGGAGCCACCTATGACGAAATCAAGGCTAAGGTCGAG
Dmela MAJ  GATCTTACCGTCCGCCTGGGCAAGGGAGCCACCTATGACGAAATCAAGGCTAAGGTCGAG
Dsimu      GATCTTACCGTCCGCCTGGGCAAGGGAGCCAGCTATGACGAAATCAAGGCCAAGGTCGAG
Dsech      GATCTTACCGTCCGCCTGGGCAAGGGAGCCAGCTATGACGAAATCAAGGCCAAGGTCGAG
Dmaur      GATCTTACCGTCCGCCTGGGCAAGGGAGCCAGCTATGACGAAATCAAGGCCAAGGTCGAG
Derec      GATCTTACTGTCCGCCTGGGCAAGGGAGCCTCCTATGACGAAATCAAGGCTAAGGTCGAG
Doren      GATCTTACTGTGCGCCTGGGCAAGGGAGCCACCTATGACGAAATCAAGGCTAAGGTCGAG
Dyaku      GACCTTACTGTCCGCCTGGGCAAGGGAGCCAGCTATGACGAAATCAAGGCTAAGGTCGAG
Dsant      GACCTTACTGTCCGCCTGGGCAAGGGAGCCAGCTATGACGAAATCAAGGCTAAGGTCGAG
```

***Gapdh1*** **SNP #9:**

```
                                                     208 T
                                                     174 C
Dmela min  GAGGCCTCCAAGGGACCCCTGAAGGGAATCCTGGGCTACACCGATGAGGAGGTGGTCTCC
Dmela MAJ  GAGGCCTCCAAGGGACCCCTGAAGGGAATCCTGGGCTACACCGACGAGGAGGTGGTCTCC
Dsimu      GAGGCCTCCAAGGGACCCCTGAAGGGAATCCTGGGCTACACCGACGAGGAGGTGGTCTCC
Dsech      GAGGCCTCCAAGGGACCCCTGAAGGGAATCCTGGGCTACACCGACGAGGAGGTGGTCTCC
Dmaur      GAGGCCTCCAAGGGACCCTTGAAGGGAATCCTGGGCTACACCGACGAGGAGGTGGTCTCC
Derec      GAGGCGTCCAAGGGACCCCTGAAGGGAATCCTGGGCTACACCGACGAGGAGGTTGTCTCC
Doren      GAGGCCTCCAAGGGACCCCTGAAGGGAATCCTGGGCTACACCGACGAGGAGGTTGTCTCC
Dyaku      GAGGCCTCCAAGGGACCCCTGAAGGGAATCCTGGGCTACACCGACGAGGAGGTGGTCTCC
Dsant      GAGGCCTCCAAGGGACCCCTGAAGGGAATCCTGGGCTACACCGACGAGGAGGTGGTCTCC
```

```
Gapdh1 SNP #10:   34 T
                 252 C
Dmela min    ACCGACTTCTTCAGCGACACCCATTCGTCTGTGTTCGACGCCAAGGCTGGCATTTCGCTG
Dmela MAJ    ACCGACTTCCTCAGCGACACCCATTCGTCTGTGTTCGACGCCAAGGCTGGCATTTCGCTG
Dsimu        ACCGACTTCCTCAGCGACACCCATTCGTCGGTGTTCGACGCCAAGGCTGGCATTTCGCTG
Dsech        ACCGACTTCCTCAGCGACACCCATTCGTCGGTGTTCGACGCCAAGGCTGGCATTTCGCTG
Dmaur        ACCGACTTCCTCAGCGACACCCATTCGTCGGTGTTCGACGCCAAGGCTGGCATTTCGCTG
Derec        ACCGACTTCCTCAGCGATACCCATTCGTCGGTCTTCGACGCCAAGGCTGGCATTTCGCTG
Doren        ACCGACTTCCTCAGCGATACCCATTCGTCGGTCTTCGACGCCAAGGCTGGCATTTCGCTG
Dyaku        ACCGACTTCCTCAGCGACACCCATTCGTCGGTCTTCGACGCCAAGGCTGGCATTTCGCTG
Dsant        ACCGACTTCCTCAGCGACACCCATTCGTCGGTCTTCGACGCCAAGGCTGGCATTTCGCTG

Dmela        AACGATAAGTTCGTCAAGCTAATCTCGTGGTACGACAACGAGTTCGGTTACTCCAACCGC
Dsimu        AACGACAAGTTCGTCAAGCTAATCTCGTGGTACGACAACGAGTTCGGTTACTCCAACCGC
Dsech        AACGACAAGTTCGTCAAGCTAATCTCGTGGTACGACAACGAGTTCGGTTACTCCAACCGC
Dmaur        AACGACAAGTTCGTTAAGCTAATCTCGTGGTACGACAACGAGTTCGGTTACTCCAACCGC
Derec        AACGACAAGTTCGTCAAGCTAATCTCGTGGTACGACAACGAGTTCGGTTACTCCAACCGC
Doren        AACGACAAATTCGTCAAGCTAATCTCGTGGTACGACAACGAGTTCGGTTACTCCAACCGC
Dyaku        AACGACAAGTTCGTCAAGCTTATCTCGTGGTACGACAACGAGTTCGGTTACTCCAACCGC
Dsant        AACGACAAGTTCGTCAAGCTTATCTCGTGGTACGACAACGAGTTCGGTTACTCCAACCGC

Dmela        GTCATCGACCTGATCAAGTATATGCAGAGCAAGGACTAA
Dsimu        GTCATCGACCTGATCAAGTATATGCAGAGCAAGGACTAA
Dsech        GTCATCGACCTGATCAAGTATATGCAGAGCAAGGACTAA
Dmaur        GTCATCGACCTGATCAAGTATATGCAGAGCAAGGACTAA
Derec        GTCATCGACCTGATCAAGTATATGCAGAGCAAGGACTAA
Doren        GTCATCGACCTGATCAAGTATATGCAGAGCAAGGACTAA
Dyaku        GTCATCGACCTCATCAAGTATATGCAGAGCAAGGACTAA
Dsant        GTCATCGACCTCATCAAGTATATGCAGAGCAAGGACTAA
```

**Figure S7. *Drosophila melanogaster Gapdh2* polymorphisms.** Shown is an alignment of *Gapdh2* and 6 different SNPs identified in DGRP data. This polymorphism data is more difficult to classify unambiguously than the *Gapdh1* data (10 SNPs) because many of the polymorphisms here are associated with imperfect TRs. Residues highlighted in cyan are derived changes. Residues highlighted in gray are unpolarized differences detected in the outgroup clade. Number of DGRP lines with unambigious genotypes are indicated.

```
Dmela        ATGTCGAAGATTGGTATCAATGGATTTGGTCGCATCGGCCGCTTGGTTCTCCGCGCCGCC
Dsimu        ATGTCGAAGATTGGTATCAATGGATTTGGTCGCATCGGCCGCTTGGTTCTGCGCGCCGCC
Dsech        ATGTCGAAGATTGGTATCAATGGATTTGGTCGCATCGGACGCTTGGTTCTGCGCGCCGCC
Dmaur        ATGTCGAAGATTGGTATCAATGGATTTGGTCGCATCGGCCGCTTGGTTCTGCGCGCCGCC
Derec        ATGTCGAAGATTGGTATCAATGGATTTGGTCGCATCGGCCGCTTGGTTCTGCGCGCCGCC
Dyaku        ATGTCGAAGATTGGTATCAACGGATTTGGCCGCATCGGTCGCTTGGTTCTGCGCGCCGCC
Dsant        ATGTCGAAGATTGGTATCAACGGATTTGGCCGCATCGGTCGCTTGGTTCTGCGCGCCGCC

Dmela        ATTGATAAGGGCGCCAACGTTGTGGCCGTCAACGATCCCTTCATCGATGTGAACTACATG
Dsimu        ATCGATAAGGGCGCCAACGTTGTGGCCGTCAACGATCCCTTCATCGATGTGAACTACATG
Dsech        ATCGATAAGGGCGCCAACGTTGTGGCCGTCAACGATCCCTTCATCGATGTGAACTACATG
Dmaur        ATCGATAAGGGCGCCAACGTTGTGGCCGTCAACGATCCCTTCATCGATGTGAACTACATG
Derec        ATCGATAAGGGTGCAAACGTTGTGGCCGTCAACGATCCCTTCATCGATGTGAACTACATG
Dyaku        ATCGATAAGGGTGCTAACGTTGTGGCCGTCAACGATCCCTTCATCGATGTGAACTACATG
Dsant        ATCGATAAGGGTGCTAACGTTGTGGCCGTCAACGATCCCTTCATCGATGTGAACTACATG

Dmela        GTCTACCTGTTCAAGTTCGATTCGACCCACGGACGTTTTAAGGGCACCGTTGCCGCCGAG
Dsimu        GTCTACCTGTTCAAGTTCGATTCGACCCACGGACGTTTTAAGGGCACCGTTGCCGCCGAG
Dsech        GTCTACCTGTTCAAGTTCGATTCGACCCACGGACGTTTTAAGGGCACCGTTGCCGCCGAG
Dmaur        GTCTACCTGTTCAAGTTCGATTCGACCCACGGACGTTTTAAGGGCACCGTTGCCGCCGAG
Derec        GTCTACCTGTTCAAGTTCGATTCGACCCACGGACGTTTTAAGGGCACCGTTGCCGCCGAG
Dyaku        GTCTACCTGTTCAAGTTCGATTCCACCCACGGACGTTTTAAGGGCACCGTTGCCGCCGAG
Dsant        GTCTACCTGTTCAAGTTCGATTCCACCCACGGACGTTTTAAGGGCACCGTTGCCGCCGAG
```

***Gapdh2* SNP #1:**

```
                                                                    76 T
                                                                   310 C
Dmela min    GGCGGTTTCCTGGTGGTCAACGGCCAGAAGATCACCGTCTTCAGCGAACGCGACCCGGTC
Dmela MAJ    GGCGGTTTCCTGGTGGTCAACGGCCAGAAGATCACCGTCTTCAGCGAACGCGACCCGGCC
Dsimu        GGCGGTTTCCTGGTGGTCAACGGCCAGAAGATCACCGTTTTCAGCGAACGCGACCCGGCC
Dsech        GGCGGTTTCCTGGTGGTCAACGGCCAGAAGATCACCGTTTTCAGCGAACGCGACCCGGCC
Dmaur        GGCGGTTTCCTGGTGGTCAACGGCCAGAAGATCACCGTTTTCAGCGAACGCGACCCGGCC
Derec        GGCGGTTTCCTGGTGGTTAACGGCCAGAAGATCACCGTCTTCAGCGAACGCGACCCGACC
Dyaku        GGCGGTTTCCTGGTGGTCAACGGCCAGAAGATCACCGTCTTCAGCGAACGCGACCCGGCC
Dsant        GGCGGTTTCCTGGTGGTCAACGGCCAGAAGATCACCGTCTTCAGCGAACGCGACCCGGCC

Dmela        AACATCAACTGGGCCAGCGCTGGTGCCGAATACATCGTGGAGTCCACTGGCGTGTTCACC
Dsimu        AACATCAACTGGGCCAGCGCTGGTGCCGAATACATCGTGGAGTCCACTGGCGTGTTCACC
Dsech        AACATCAACTGGGCCAGCGCTGGTGCCGAATACATCGTGGAGTCCACTGGCGTGTTCACC
Dmaur        AACATCAACTGGGCCAGCGCTGGTGCCGAATATATCGTGGAGTCCACTGGCGTGTTCACC
Derec        AACATCAACTGGGCCAGCGCTGGTGCCGAATATATCGTGGAGTCCACTGGCGTGTTCACC
Dyaku        AACATCAACTGGGCCAGCGCTGGTGCCGAATACATCGTGGAGTCCACTGGTGTGTTCACC
Dsant        AACATCAACTGGGCCAGCGCTGGTGCCGAATACATCGTGGAGTCCACTGGTGTGTTCACC
```

```
Dmela      ACCATCGACAAGGCATCCACTCACTTGAAGGGCGGTGCCAAGAAGGTTATCATCTCGGCC
Dsimu      ACCATCGACAAGGCATCCACTCACTTGAAGGGCGGTGCCAAGAAGGTTATCATCTCGGCC
Dsech      ACCATCGACAAGGCATCCACTCACTTGAAGGGCGGTGCCAAGAAGGTTATCATCTCGGCC
Dmaur      ACCATCGACAAGGCCTCCACTCACTTGAAGGGCGGTGCCAAGAAGGTTATCATCTCGGCC
Derec      ACCATCGACAAGGCATCCACTCACTTGAAGGGCGGTGCCAAGAAGGTGATCATCTCGGCC
Dyaku      ACCATCGACAAGGCATCCACTCACTTGAAGGGCGGTGCCAAGAAGGTGATCATCTCGGCC
Dsant      ACCATCGACAAGGCATCCACTCACTTGAAGGGCGGTGCCAAGAAGGTGATCATCTCGGCC

Dmela      CCATCCGCCGATGCTCCCATGTTCGTTTGCGGCGTCAACTTGGATGCCTACAAGCCCGAC
Dsimu      CCATCCGCCGATGCTCCCATGTTCGTTTGCGGCGTCAACTTGGATGCCTACAAGCCCGAC
Dsech      CCATCCGCCGATGCTCCCATGTTCGTTTGCGGTGTCAACTTGGATGCCTACAAGCCCGAC
Dmaur      CCATCCGCCGATGCTCCCATGTTCGTTTGCGGTGTCAACTTGGATGCCTACAAGCCCGAC
Derec      CCATCCGCTGATGCTCCCATGTTCGTCTGCGGCGTCAACTTGGATGCCTACAATCCGGAC
Dyaku      CCATCCGCCGATGCTCCCATGTTCGTGTGCGGCGTCAACTTGGATGCCTACAAGCCGGAC
Dsant      CCATCTGCCGATGCTCCCATGTTCGTCTGCGGCGTCAACTTGGATGCCTACAAGCCGGAC

Dmela      ATGAAGGTGGTCTCCAACGCATCGTGCACCACCAACTGCTTGGCTCCTCTGGCCAAGGTG
Dsimu      ATGAAGGTGGTCTCCAACGCATCGTGCACCACCAACTGCCTGGCTCCTCTGGCCAAGGTG
Dsech      ATGAAGGTGGTCTCCAACGCATCGTGCACCACCAACTGCCTGGCTCCTCTGGCCAAGGTG
Dmaur      ATGAAGGTGGTCTCCAACGCATCGTGCACCACCAACTGCCTGGCTCCTCTGGCCAAGGTG
Derec      ATGAAGGTGGTTTCCAACGCATCGTGCACCACCAACTGCCTGGCTCCTTTGGCCAAGGTG
Dyaku      ATGAAGGTGGTTTCCAACGCATCGTGCACCACCAACTGCCTGGCTCCATTGGCCAAGGTG
Dsant      ATGAAGGTGGTTTCCAACGCATCGTGCACCACCAACTGCCTGGCTCCATTGGCCAAGGTG

Dmela      ATCAACGACAACTTCGAGATCGTCGAGGGTCTGATGACCACCGTTCATGCCACCACCGCT
Dsimu      ATCAACGACAACTTCGAGATCGTCGAGGGTCTGATGACCACCGTTCATGCCACCACTGCT
Dsech      ATCAACGACAACTTCGAGATCGTCGAGGGTCTGATGACCACCGTTCATGCCACCACCGCT
Dmaur      ATCAACGACAACTTCGAGATCGTCGAGGGTCTGATGACCACCGTTCATGCCACCACCGCT
Derec      ATCAACGATAACTTCGAAATCGTCGAGGGTCTGATGACCACCGTTCATGCCACCACCGCT
Dyaku      ATCAACGACAACTTCGAGATCGTCGAGGGTCTGATGACCACCGTTCATGCCACCACCGCT
Dsant      ATCAACGACAACTTCGAGATCGTCGAGGGTCTGATGACCACCGTTCATGCCACCACCGCT

Dmela      ACCCAGAAGACCGTCGATGGACCTTCCGGCAAGTTGTGGCGTGATGGACGTGGCGCTGCC
Dsimu      ACCCAGAAGACCGTCGATGGACCTTCCGGCAAGTTGTGGCGTGATGGACGTGGCGCTGCC
Dsech      ACCCAGAAGACCGTCGATGGACCTTCCGGCAAGTTGTGGCGTGATGGACGTGGCGCTGCC
Dmaur      ACCCAGAAGACCGTCGATGGACCTTCCGGCAAGTTGTGGCGTGATGGACGTGGCGCTGCC
Derec      ACCCAGAAGACCGTCGATGGACCGTCCGGCAAGTTGTGGCGTGATGGACGTGGCGCTGCC
Dyaku      ACCCAGAAGACCGTCGATGGACCTTCCGGCAAGTTGTGGCGTGATGGACGTGGCGCTGCC
Dsant      ACCCAGAAGACCGTCGATGGACCTTCCGGCAAGTTGTGGCGTGATGGACGTGGCGCTGCC

Dmela      CAGAACATCATTCCAGCTTCCACTGGAGCTGCCAAGGCCGTGGGCAAGGTTATCCCCGCC
Dsimu      CAGAACATCATTCCAGCTTCCACCGGAGCTGCCAAGGCCGTGGGCAAGGTTATCCCCGCC
Dsech      CAGAACATCATTCCAGCTTCCACCGGAGCTGCCAAGGCCGTGGGCAAGGTTATCCCCGCC
Dmaur      CAGAACATCATTCCAGCTTCCACCGGAGCTGCCAAGGCCGTGGGCAAGGTTATCCCCGCC
Derec      CAGAACATCATTCCAGCTTCGACCGGAGCTGCCAAGGCCGTGGGCAAGGTTATTCCCGCC
Dyaku      CAGAACATCATTCCAGCTTCCACCGGAGCTGCCAAGGCCGTGGGCAAGGTTATCCCCGCC
Dsant      CAGAACATCATTCCAGCTTCCACCGGAGCTGCCAAGGCCGTGGGCAAGGTTATCCCCGCC
```

```
Gapdh2 SNP #2:                                    2 A
                                                408 G
Dmela min    CTCAACGGTAAGCTCACCGGAATGGCATTCCGTGTACCCACTCCCAACGTTTCCGTGGTC
Dmela MAJ    CTCAACGGTAAGCTCACCGGAATGGCATTCCGTGTGCCCACTCCCAACGTTTCCGTGGTC
Dsimu        CTCAATGGTAAGCTCACCGGAATGGCATTCCGTGTGCCCACTCCCAACGTTTCCGTGGTC
Dsech        CTCAATGGTAAGCTCACCGGAATGGCATTCCGTGTGCCCACTGCCAACGTTTCCGTGGTC
Dmaur        CTCAATGGTAAGCTCACCGGAATGGCATTCCGTGTGCCCACTCCCAACGTTTCCGTGGTC
Derec        CTCAACGGTAAGCTCACCGGAATGGCATTCCGTGTGCCCACTCCCAACGTTTCCGTGGTC
Dyaku        CTCAATGGTAAGCTCACCGGAATGGCATTCCGTGTGCCCACTCCCAACGTTTCCGTGGTC
Dsant        CTCAATGGTAAGCTCACCGGAATGGCATTCCGTGTGCCCACTCCCAACGTTTCCGTGGTC

Dmela        GATTTGACCGTGCGCTTGGGCAAGGGTGCGTCCTATGATGAAATTAAGGCCAAGGTTCAG
Dsimu        GATTTGACCGTGCGCTTGGGCAAGGGTGCGTCCTATGATGAAATTAAGGCCAAGGTTCAG
Dsech        GATTTGACCGTGCGCTTGGGCAAGGGTGCGTCCTATGATGAAATTAAGGCCAAGGTTGAG
Dmaur        GATTTGACCGTGCGCTTGGGCAAGGGTGCGTCCTATGATGAAATTAAGGCCAAGGTTCAG
Derec        GATCTGACCGTGCGCTTGGGCAAGGGTGCATCCTATGATGAAATTAAGGCCAAGGTTCAG
Dyaku        GATTTGACCGTGCGCTTGGGCAAGGGTGCATCCTACGATGAAATTAAGGCCAAGGTACAG
Dsant        GATTTGACCGTGCGCTTGGGCAAGGGTGCATCCTACGATGAAATTAAGGCCAAGGTACAG
```

```
Gapdh2 SNPs #3-4:                           2 T                  2 A
                                          408 C                408 G
Dmela min    GAGGCCGCCAACGGACCCCTGAAGGGTATTCTGGGATACACCGATGAGGAAGTCGTTTCT
Dmela MAJ    GAGGCCGCCAACGGACCCCTGAAGGGTATCCTGGGATACACCGATGAGGAGGTCGTTTCT
Dsimu        GAGGCCGCCAACGGACCCCTGAAGGGAATCCTGGGATACACCGATGAGGAGGTCGTGTCC
Dsech        GAGGCCGCCAACGGACCCCTGAAGGGAATCCTGGGATACACCGATGAGGAGGTCGTGTCC
Dmaur        GAGGCCGCCAACGGACCCCTGAAGGGAATCCTGGGATACACCGATGAGGAGGTCGTGTCC
Derec        GAGGCCGCCAACGGACCCCTGAAGGGAATTTTGGGATACACCGATGAGGAGGTCGTGTCC
Dyaku        GAGGCCGCCAACGGACCCCTGAAGGGAATCTTGGGATACACCGATGAGGAGGTCGTGTCC
Dsant        GAGGCCGCCAACGGACCCCTGAAGGGAATCTTGGGATACACCGATGAGGAGGTCGTGTCC
```

```
Gapdh2 SNP #5:                                        2 T
                                                    406 G
Dmela min    ACCGATTTCCTCAGCGACACCCACTCGTCGGTGTTCGATTCCAAGGCTGGCATTTCGCTA
Dmela MAJ    ACCGATTTCCTCAGCGACACCCACTCGTCGGTGTTCGATGCCAAGGCTGGCATTTCGCTA
Dsimu        ACCGATTTCCTCAGCGACACCCACTCGTCGGTGTTCGATGCCAAGGCTGGCATTTCGCTG
Dsech        ACCGATTTCCTCAGCGACACCCACTCGTCGGTGTTCGATGCCAAGGCTGGCATTTCGCTG
Dmaur        ACCGATTTCCTCAGCGACACCCACTCGTCGGTGTTCGATGCCAAGGCTGGCATTTCGCTG
Derec        ACCGATTTCCTCAGCGACACCCACTCGTCGGTCTTCGATGCCAAGGCTGGCATTTCGCTG
Dyaku        ACCGATTTCCTCAGCGACACCCACTCGTCGGTGTTCGATGCCAAGGCTGGCATTTCGCTG
Dsant        ACCGATTTCCTCAGCGACACCCACTCGTCGGTGTTCGATGCCAAGGCTGGCATTTCGCTG
```

```
Gapdh2 SNP #6:      4 T
                  400 C
Dmela min    AACGACAAGTTTGTGAAGCTGATCTCTTGGTACGACAACGAGTTTGGCTACTCCAACCGC
Dmela MAJ    AACGACAAGTTCGTGAAGCTGATCTCTTGGTACGACAACGAGTTTGGCTACTCCAACCGC
Dsimu        AACGACAAGTTCGTGAAGCTGATCTCTTGGTACGACAACGAGTTTGGATACTCCAACCGC
Dsech        AACGACAAGTTCGTGAAGCTGATCTCTTGGTACGACAACGAGTTTGGATACTCCAACCGC
Dmaur        AACGACAAGTTCGTGAAGCTGATCTCTTGGTACGACAACGAGTTTGGATACTCCAACCGC
Derec        AACGACAAGTTCGTGAAGCTGATCTCTTGGTACGACAACGAGTTTGGATACTCCAACCGC
Dyaku        AACGACAAGTTCGTGAAGCTGATCTCTTGGTACGACAACGAGTTTGGATACTCCAACCGC
Dsant        AACGACAAGTTCGTGAAGCTGATCTCTTGGTACGACAACGAGTTTGGATACTCCAACCGC
```

```
Dmela      GTCATCGACCTGATCAAGTACATGCAGAGCAAGGATTAA
Dsimu      GTCATCGACCTGATCAAGTACATGCAGAGCAAGGATTAA
Dsech      GTCATCGACCTGATCAAGTACATGCAGAGCAAGGATTAA
Dmaur      GTCATCGACCTGATCAAGTACATGCAGAGCAAGGATTAA
Derec      GTCATCGATCTGATCAAGTACATGCAGAGCAAGGATTAA
Dyaku      GTCATCGACCTGATCAAGTACATGCAGAGCAAGGATTAA
Dsant      GTCATCGACCTGATCAAGTACATGCAGAGCAAGGATTAA
```

**Table S1. TR content within the highly conserved *gapdh* CDS is substantial but fast evolving.** Repeat content for the *gapdh* CDS is shown for seven different insect species. None of this repeat content is shared across all seven lineages. A comparison between *gapdh* sequences from *Drosophila melanogaster* and the mosquito *Anopheles gambiae* shows that ~50% of repeat content in each lineage is unique to that lineage. Numbers in parentheses indicate the number of repeats unique to each species in the pairwise comparison of *LINEUP* data (see also Fig. S1).

| Gene | Total | *k*=2 | *k*=3 | *k*=4 | *k*=5 | *k*=6 | *k*=7 | *k*=8 |
|---|---|---|---|---|---|---|---|---|
| *D. mela. gapdh1* | 55 | 26 (13) | 27 (12) | 1 (1) | 1 (1) | 0 (0) | 0 (0) | 0 (0) |
| *D. mela. gapdh2* | 46 | 22 | 22 | 1 | 1 | 0 | 0 | 0 |
| *M. dome. gapdh* | 34 | 18 | 15 | 1 | 0 | 0 | 0 | 0 |
| *S. calc. gapdh* | 41 | 25 | 14 | 1 | 1 | 0 | 0 | 0 |
| *A. gamb. gapdh* | 53 | 25 (14) | 27 (13) | 0 (0) | 0 (0) | 0 (0) | 0 (0) | 1 (1) |
| *B. mori gapdh* | 46 | 28 | 16 | 2 | 0 | 0 | 0 | 0 |
| *A. glab. gapdh* | 38 | 19 | 17 | 3 | 0 | 0 | 0 | 0 |
| Shared (all genes) | 0 | 0 | 0 | 0 | 0 | 0 | 0 | 0 |

**Table S2. Trinucleotide repeats occur randomly across all three reading frames of the *gapdh* CDS.** Due to the high turnover of TR content, most TRs are lineage-specific between divergent lineages, thus allowing meaningful comparisons of independent mutational histories for the same gene. Here, I show the proportion of triplet repeats in the *gapdh* CDS of *D. melanogaster*, *A. gambiae*, and the moth *Bombyx mori* (999 bp each). Trinucleotide repeats occur in any frame, and anywhere from 1/3 to 2/3 of all trinucleotide repeats from one lineage do not occur in the codon frame. A null model of trinucleotide repeat origination would predict 1/3 of single frame TRs to happen in the codon frame. This is roughly what is seen (last column). Altogether, these results suggest that trinucleotide repeats are not the result of positive selection for repeated amino acids but rather the result of the mutational force of replication slippage. Importantly, none of these trinucleotide repeats produce indels at the amino acid level. Instead, these trinucleotide repeats replace adjacent sequences, often resulting in multiple apparent transversion-rich substitutions.

| Gene | # trinucl. TRs | NOT in codon frame in any cycling frame | one frame only (no cycling) | single frame = codon frame |
|---|---|---|---|---|
| *D. mela. gapdh1* | 27 | 0.370 (10/27) | 0.556 (15/27) | 0.333 ( 5/15) |
| *A. gamb. gapdh* | 27 | 0.481 (13/27) | 0.667 (18/27) | 0.278 ( 5/18) |
| *B. mori gapdh* | 18 | 0.750 (12/16) | 0.812 (13/16) | 0.154 ( 2/13) |
| **Total** | **72** | **0.500 (35/70)** | **0.657 (46/70)** | **0.261 (12/46)** |
